# A Universal RT-qPCR Method for DNA Aptamer Quantification

**DOI:** 10.1101/2025.06.30.662296

**Authors:** Lijing Wang, Anran Liu, Kexin Li, Xiujuan Lv, Meng Lin, Hui Fu, Liang Hu

## Abstract

Aptamers are widely used in various applications; however, their quantification methods remain underdeveloped. In this study, we established a universal RT-qPCR-based method for DNA aptamer quantification. By incorporating reverse transcription, primers and probes could be added to aptamers, enabling their detection via qPCR (Figure 1). We developed a software tool for primer and probe design and optimized the reverse transcription process. Two aptamers, CD9-aptamer and CD63-aptamer, were selected as model systems representing two distinct aptamer types: the CD9-aptamer contains an intrinsic primer-binding site within its sequence, whereas the CD63-aptamer does not. Using this method, the limit of detection (LOD) for CD9-aptamer reached 10^-14^ M, while the LOD for CD63-aptamer was 10^-12^ M. Compared to existing quantification methods, this approach significantly improves accuracy and cost-effectiveness. Additionally, the method supports SYBR Green, TaqMan, and one-step RT-qPCR assays, broadening its applicability and enhancing precision for various aptamer-based applications.

## Introduction

Aptamers are short, single-stranded nucleic acids (DNA or RNA) that can bind to specific target molecules with high affinity and specificity. They are generated through an in vitro selection process known as Systematic Evolution of Ligands by EXponential Enrichment (SELEX), which iteratively screens large nucleic acid libraries to identify sequences that strongly interact with their target. Aptamers exhibit several unique advantages over traditional recognition molecules such as antibodies. They can be synthesized chemically, ensuring high reproducibility and cost-effectiveness. Unlike antibodies, aptamers generally exhibit less of an immune response and can be modified to enhance their stability against enzymatic degradation. Their small size allows for better tissue penetration, making them highly versatile in biomedical applications. Due to their remarkable binding properties, aptamers have been widely used in therapeutics, diagnostics, biosensing, and targeted drug delivery. They are employed in cancer therapy, viral detection, and biomarker identification, demonstrating their potential as powerful tools in modern medicine and biotechnology.^1^

Despite their promising applications, the use of aptamers faces several challenges, particularly in the area of quantification. Quantification of aptamers is crucial for its applications, such as aptamer drug pharmacokinetics^2^ and aptamer-based quantification methods like Enzyme-Linked Aptamer Sorbent Assay (ELASA)^3^. According to the aptamer database (http://:www.aptamer.org/aptamerdb)” www.aptamer.org/aptamerdb), as of January 2021, a total of 4,178 aptamers have been reported, including 3,029 DNA aptamers. While RNA aptamers are typically quantified using reverse transcription polymerase chain reaction (RT-PCR)^4,5^, no simple and cost-effective method exists for quantifying DNA aptamers. Existing approaches, such as high-performance liquid chromatography (HPLC)^2^ or isotope labeling^6^, are expensive, cumbersome and have relatively low accuracy.

Ideally, DNA aptamers should be quantifiable through quantitative PCR (qPCR) without requiring structural modifications. However, previous attempts have faced limitations. Few aptamers are long enough to allow the use of standard primer design tools.^7,8^ Katherine Perschbacher et al.^9^ developed long primers (40 bp) with a hairpin structure for qPCR, but this approach may not be universally applicable to all aptamer sequences and still resulted in primer dimers in our tests. Zaizai Dong et al.^10^ ligated primers to both ends of the aptamer, but the addition of extra sequences could potentially interfere with aptamer-target binding^11^.

In this study, we developed a reverse transcription-quantitative PCR (RT-qPCR) method to quantify aptamers without requiring structural modifications. A custom software tool was created to design the primers and probes, and reverse transcription was used to incorporate these elements, enabling aptamer quantification through RT-qPCR (Figure 1).

**Figure 1.**
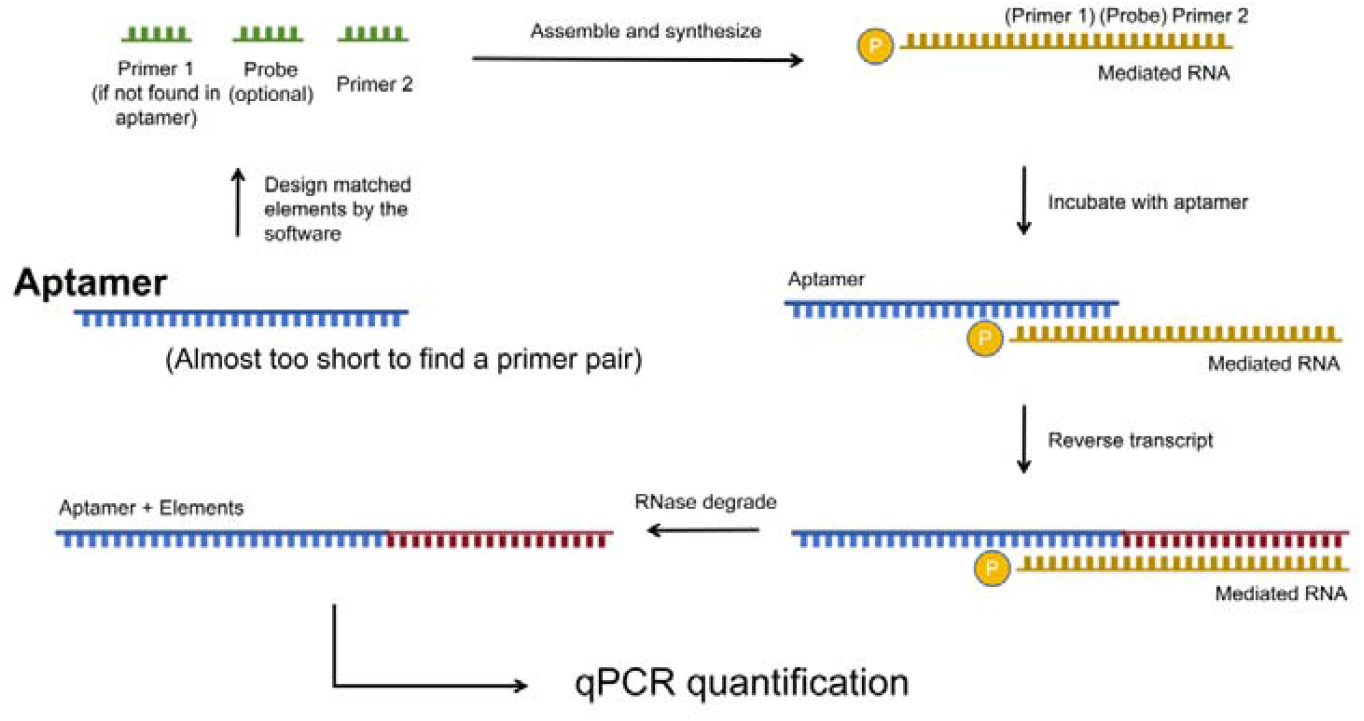
Schematic illustration of RT-qPCR-Based Aptamer Quantification.

## Methods

### 1. Materials

Primers and DNA sequences were synthesized by Tsingke Biotechnology Co., Ltd. (China). RNA sequences were synthesized by GENCEFE Biotech (Wuxi, China). AMV Reverse Transcriptase (BBI Solutions), Tth DNA Polymerase (Yeasen) were purchased for use in this study. qPCR was performed using Thermo Fisher’s QuantStudio 5 or 6.

### 2. Generation of Primers, Probes, and Mediated RNA

Aptamers can be broadly classified into three categories based on the presence of primer-binding sites: (i) two primer sites, (ii) one primer site, or (iii) no primer site. For aptamers containing two primer sites, conventional online DNA primer design tools are typically sufficient. In contrast, aptamers with only one or no primer site require the introduction of additional sequence elements designed by the software, instructions for which can be found in the Supplementary Materials 2. The user interface (UI) of the software is shown in Figure 2, and the main process is illustrated in Figure 3.

**Figure 2.**
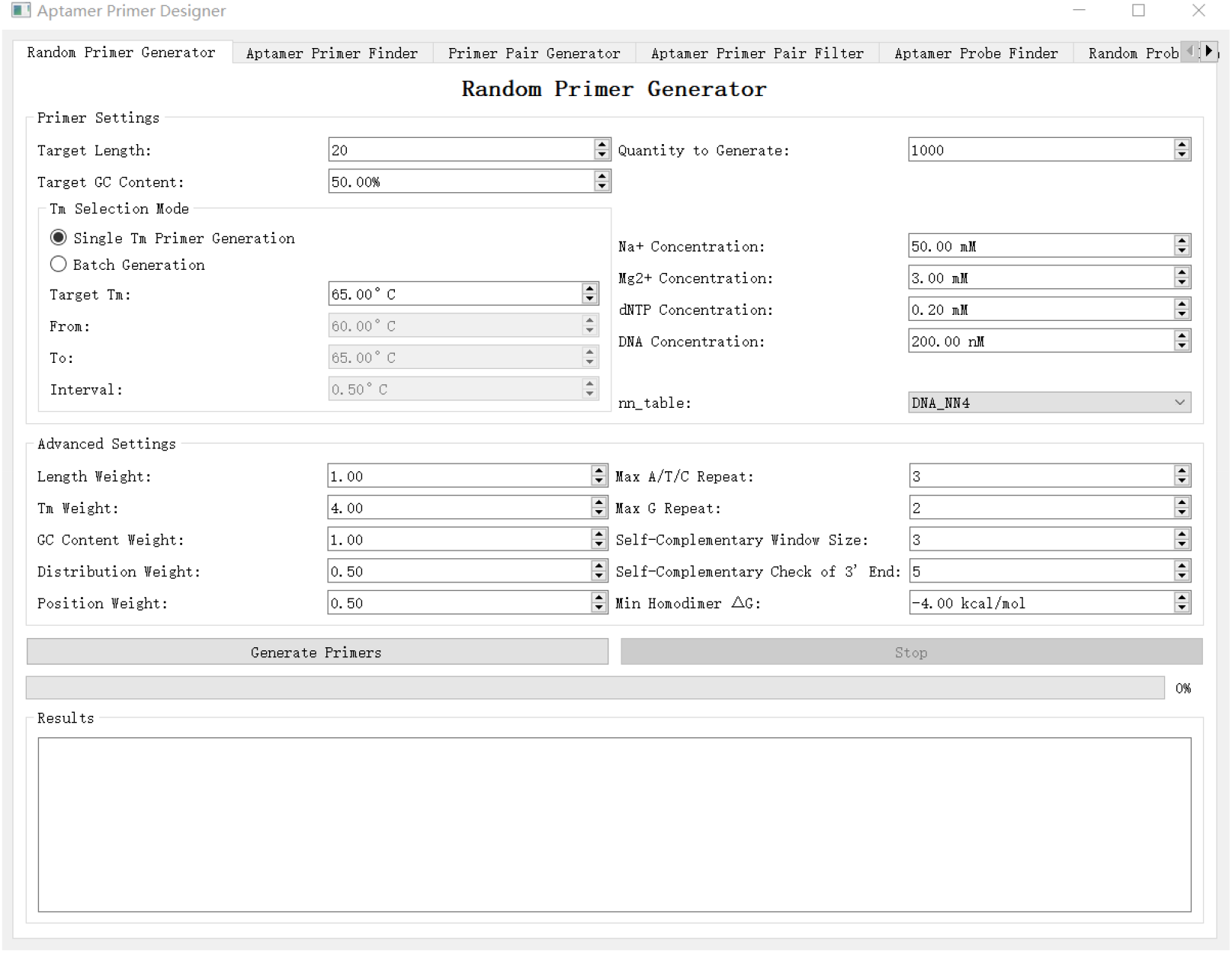
UI of the “Aptamer Primer Designer” software.

**Figure 3.**
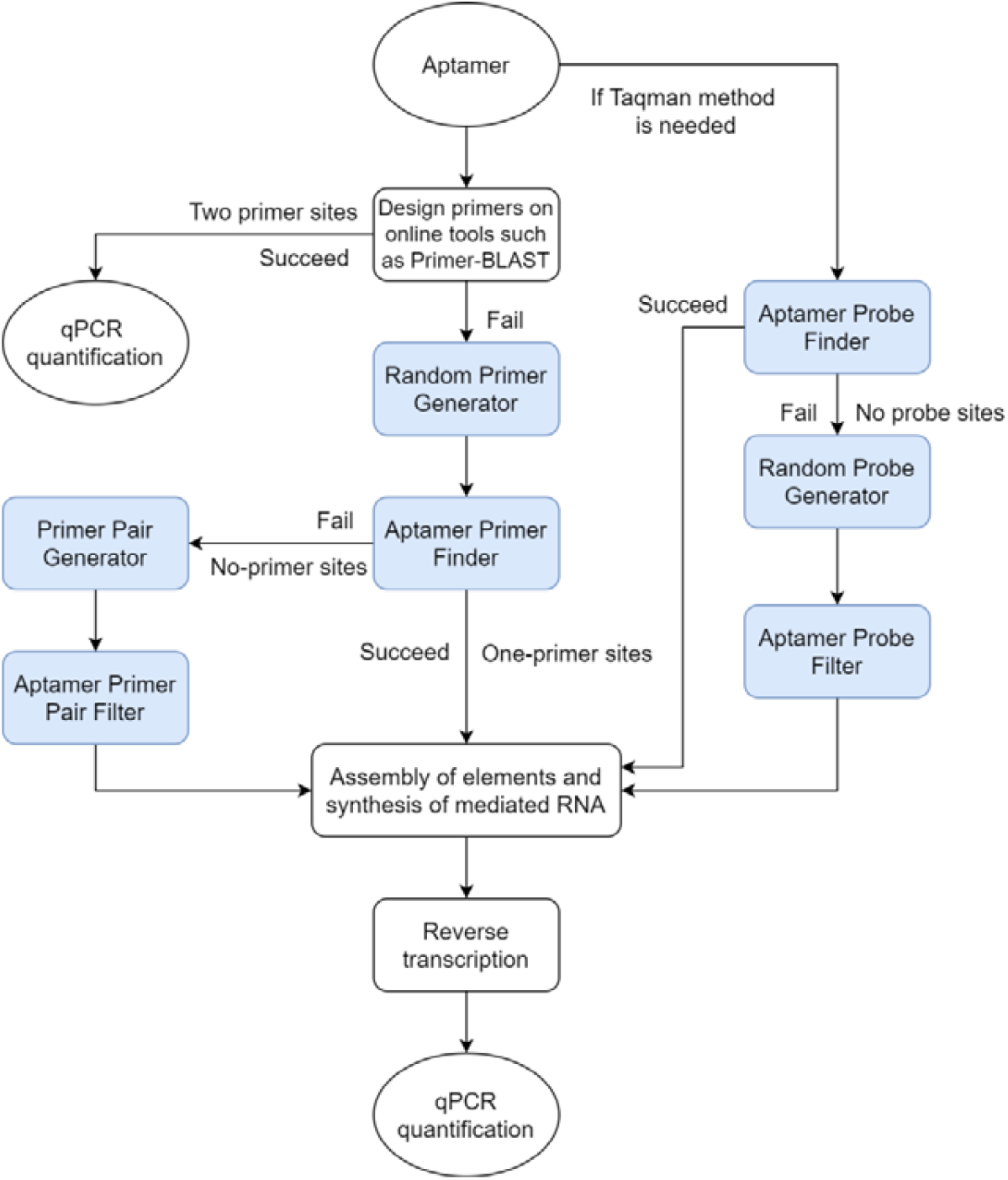
Flowchart for using “Aptamer Primer Designer” software.

The basic principle is to generate a large number of random primers and then screen for those that are compatible with the aptamer sequence.

#### Random Primer Generation

Initially, random sequences with varying melting temperatures (Tm) were produced by the “Random Primer Generator,” which relied on a random mutation algorithm constrained by standard primer design requirements.^12,13^ These sequences served as a resource for subsequent steps.

#### Primer Selection for Aptamer

Next, the “Aptamer Primer Finder” algorithm identified potential primers within each aptamer and paired them with the random primer pool to form functional primer sets. If no suitable primer was detected, a new primer-pair pool was generated with the “Primer Pair Generator,” and viable pairs were subsequently selected using the “Aptamer Primer Pair Filter.” It was recommended to use Primer-BLAST to verify whether the primer interacts with the target species.

#### Probe Design for TaqMan Assays

In addition to SYBR-based qPCR, the workflow also supports TaqMan assays. In such cases, the “Apt Probe Finder” checked whether a suitable probe could be located directly within the aptamer sequence. If no appropriate probe was identified, the “Random Probe Generator” created a probe pool, from which the “Apt Probe Filter” then selected valid probe candidates.

#### Terminal Sequence Selection and Mediated RNA Synthesis

Finally, a suitable segment was identified at the aptamer terminus to ensure that the mediated RNA could bind effectively. This segment was designed to have a melting temperature above the reverse transcription reaction temperature (50°c in our protocol) but was not so long as to induce nonspecific interactions (a Tm of 55°cwas available in this study). Once this terminal sequence was determined—together with the necessary probes and primers—these elements were assembled, and the mediated RNA was synthesized for subsequent use in the reverse transcription.

#### Implementation

All modules were written in Python and utilized the “Biopython”, “ViennaRNA” and “Primer3” packages.^14-16^

### 3. Reverse transcription

The amount of mediated RNA can be adjusted, but it should exceed the amount of aptamer in the final sample. In this study, 1.5 uL of 10^-7^ M mediated RNA was used in a 20 uL reaction. Initially, the mediated RNA and x (<=12.2) uL of the aptamer sample were added and heated to 75+°c for 5 minutes in a water bath and then cooled to reset the structure. Than, 4 uL of 5X Reaction Buffer of reverse transcription and 0.5ul RNase R (diluted 1:100 prior to use; G3462, Servicebio) were added. After 20 min reaction in 50°c bath, 1 uL of BeyoRT™ III M-MLV reverse transcriptase (D7176, Beyotime), 0.8 uL of dNTP Mix (25 mM, Beyotime), and (12.2-x) uL of ddH_2_O were mixed and added to the tube cap. Before starting the reaction, the components were mixed by slight centrifugation to bring the contents together for reducing nonspecific transcription at room temperature. The reaction was then incubated at 50°c for 20 minutes for reverse transcription and at 80°c for 10 minutes to inactivate the transcriptase.

Optional: to degrade the mediated RNA, 1 uL of 10 mg/mL RNase A (Phygene, China) was added to each sample, followed by incubation at 65°c for 10 minutes.

It should be noted that for standard curves and samples, especially those at low concentrations, a nucleic acid carrier (such as another aptamer or salmon sperm DNA) should be added to improve repeatability. In this study, CD9-aptamer standard curve samples were diluted using a 10^-10^ M CD63-aptamer solution.

### 4. qPCR

For the SYBR-based method, the following components were mixed for each sample (final volume 20ul): 2 uL of reverse transcription product, 0.4 uL of 10 uM forward primer (F), 0.4 uL of 10 uM reverse primer (R), 10 uL of ChamQ Universal SYBR qPCR Master Mix (Vazyme), and 7.2 uL of ddH_2_O.

For the TaqMan-based method, the components for each sample included (final volume 20ul): 2 uL of reverse transcription product, 0.4 uL of 10 uM forward primer (F), 0.4 uL of 10 uM reverse primer (R), 0.2 uL of 10 uM probe, 10 uL of AceQ® Universal U+ Probe Master Mix V2 (Vazyme), and 7 uL of ddH_2_O.

The qPCR procedure included an initial denaturation at 95°c for 5 minutes, followed by 50 cycles of 95°c for 5 seconds and 60°c for 30 seconds.

### 5. One-step RT-qPCR

For a 20 uL reaction, 1.5 uL of 10^-7^ M mediated RNA and x (<=7.7) uL of aptamer sample were mixed, heated to 75°C for 5 min, and then cooled. The UniPeak U+ One-Step RT-qPCR SYBR Green Kit (Q226, Vazyme) was used according to the manufacturer’s instructions. Specifically, each reaction contained 10 uL of One-Step mix, 0.4 uL of each primer (10 uM), 1.5 + x uL of sample mixture, and 7.7 − x uL ddH_2_O. The RT-qPCR cycling conditions were as follows: reverse transcription at 50°C for 10 min, initial denaturation at 95°C for 30 s, followed by 50 cycles of 95°C for 10 s and 60°C for 30 s.

## Results

### 1. Design and Selection of CD9 and CD63 Aptamer Primers, Probes, and Mediated RNA

Primers for CD9 aptamer^17^ and CD63 aptamer^18^, along with probes, were designed using the “Aptamer Primer Designer” software. To create primer pools, random primers with Tm ranging from 60°c to 65°c were generated by “Random Primer Generator”. For the CD9-aptamer, primer sites were identified using the “Apt Primer Finder” and then matched with the random primer pools to form primer pairs. Among the many potential primer pairs, six (Supplementary 1 Table 1) were ramdomly selected for qPCR analysis. Three of these pairs were chosen based on the absence of amplification curves in template-free reactions, confirming that no primer dimers were formed. One pair was then selected for further steps (Table 1).

**Table 1.**
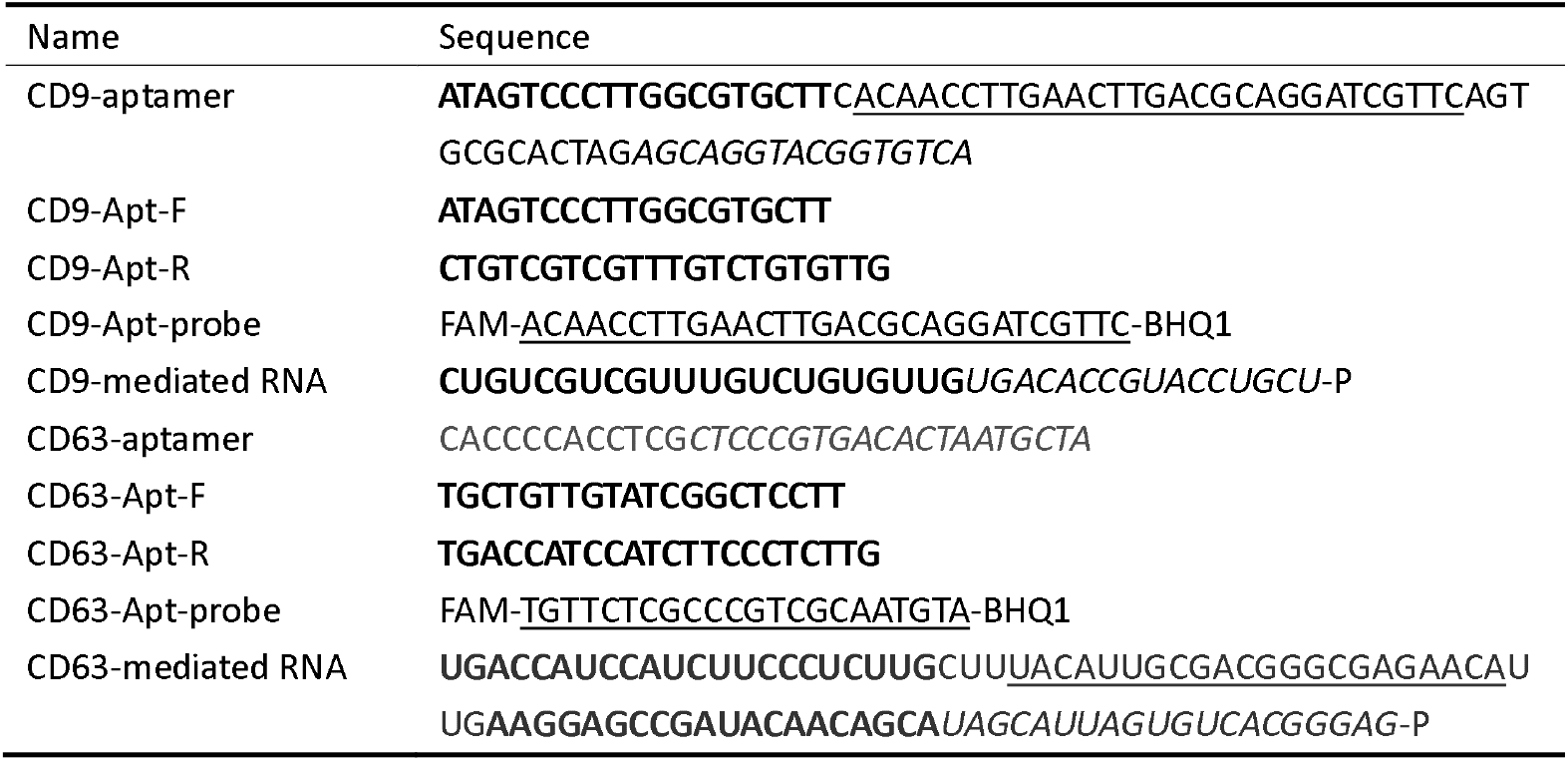
Sequences of Aptamers, Primers, Probes and Mediated RNAs designed by the software.

For CD63-aptamer, no suitable primer sites were found in the sequence. Consequently, primer pairs were generated from random primer pools using the “Primer Pair Generator.” These were then screened with the “Aptamer Primer Pair Filter” to identify optimal pairs. Three pairs exhibiting minimal secondary structure were selected (Supplementary 1 Table 2) and tested by qPCR, with two passing the test. One of these was chosen for subsequent use (Table 1).

In cases where the TaqMan method is required, probe sequences can either be identified or synthesized. Probe sequences for CD9-aptamer were identified using the “Aptamer Probe Finder,” which directly located a suitable probe within the aptamer sequence. However, no suitable probe for CD63-aptamer could be found using this method, so an alternative approach was adopted. The “Random Probe Generator” was used to create probe pools for CD63-aptamer, and these were filtered using the “Aptamer Primer Probe Filter” to minimize mismatch and secondary structure (Table 1).

The terminal regions of the CD9-aptamer and CD63-aptamer were selected based on a target Tm of approximately 60°c. All elements were assembled, and the mediated RNA was synthesized.

### 2. Optimization of Reverse Transcription

#### Concentration Comparison of Mediated RNA

The effects of different concentrations of CD9-aptamer mediated RNA (10^-5^, 10^-6^, 10^-7^, 10^-8^ M) at a volume of 1.5 uL were evaluated in the qPCR assay. The standard amplification curve for the 10^-5^ M group (Figure 4A) exhibited irregularities, suggesting that over-concentration of the mediated RNA interferes with qPCR amplification. In contrast, the curves for the 10^-6^, 10^-7^, and 10^-8^ M concentrations (Figure 4A) demonstrated a linear relationship.

**Figure 4.**
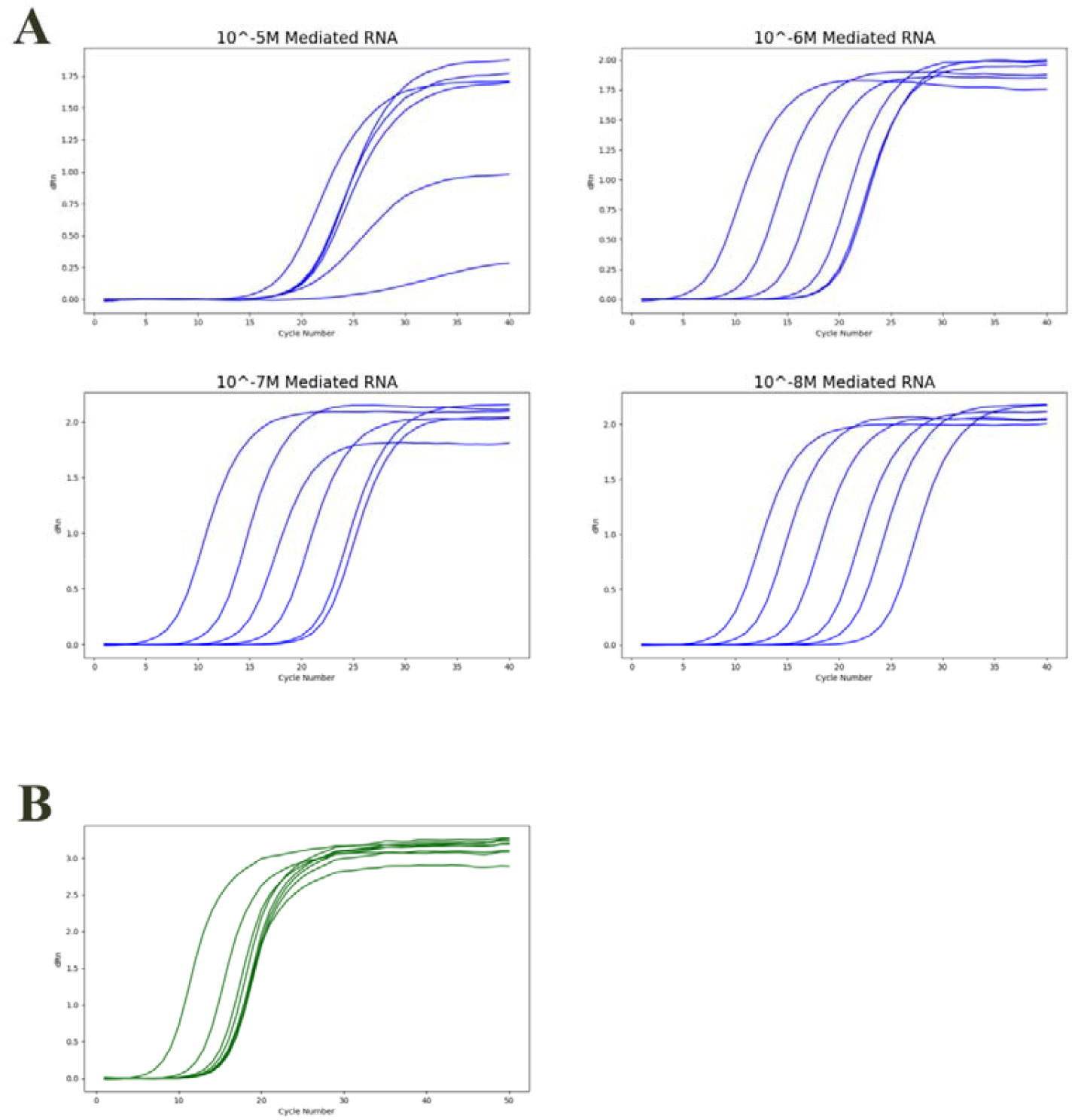
Standard Amplification Plots of RT-qPCR-Based Aptamer Quantification. (A) Amplification plots of CD9-aptamer at concentrations 10^-8^, 10^-9^, …, 10^-13^ M, with varying mediated RNA concentrations in RT (10^-5^, 10^-6^, 10^-7^, 10^-8^ M). (B) Amplification plots of CD63-aptamer at concentrations 10^-8^, 10^-9^, …, 10^-13^ M, with 10^-7^ M mediated RNA.

Then, we tested 10^-7^ M CD63-aptamer mediated RNA to quantify CD63-aptamer. However, the amplification curves appeared abnormal (Figure 4B). We proposed two possible explanations. First, non-specific products might have been generated during reverse transcription due to RNA self-priming. Second, the mediated RNA itself may have undergone amplification during the qPCR process.

#### Phosphorylation of the 3’ End of Mediated RNA Reduces Abnormal Amplification of CD-63Aptamer in qPCR Due to RNA Self-Priming

To prevent RNA self-priming, we phosphorylated the 3’ terminal of the mediated RNA. While qPCR results showed that this modification partially alleviated the issue, it did not completely resolve it (Figure 5A), indicating that some undesired DNA sequences were still generated during reverse transcription. Additional two sequences were also tested, and amplification curves were observed in no-primer reverse transcription reactions (Figure 5B, Supplementary 1 Table 3). Based on these observations, we hypothesized that the phosphorylation might be too small to fully block reverse transcription. Although reverse transcription might not initiate directly from the phosphorylated terminal, reverse transcriptase could stall at this site, allowing a random nucleotide to bind to the adjacent base and potentially trigger reverse transcription. However, despite testing other modifications, such as poly-U and C18 with steric hindrance, no improvement was observed (data not shown). This suggests that reverse transcription may occur even in the absence of a primer.

**Figure 5.**
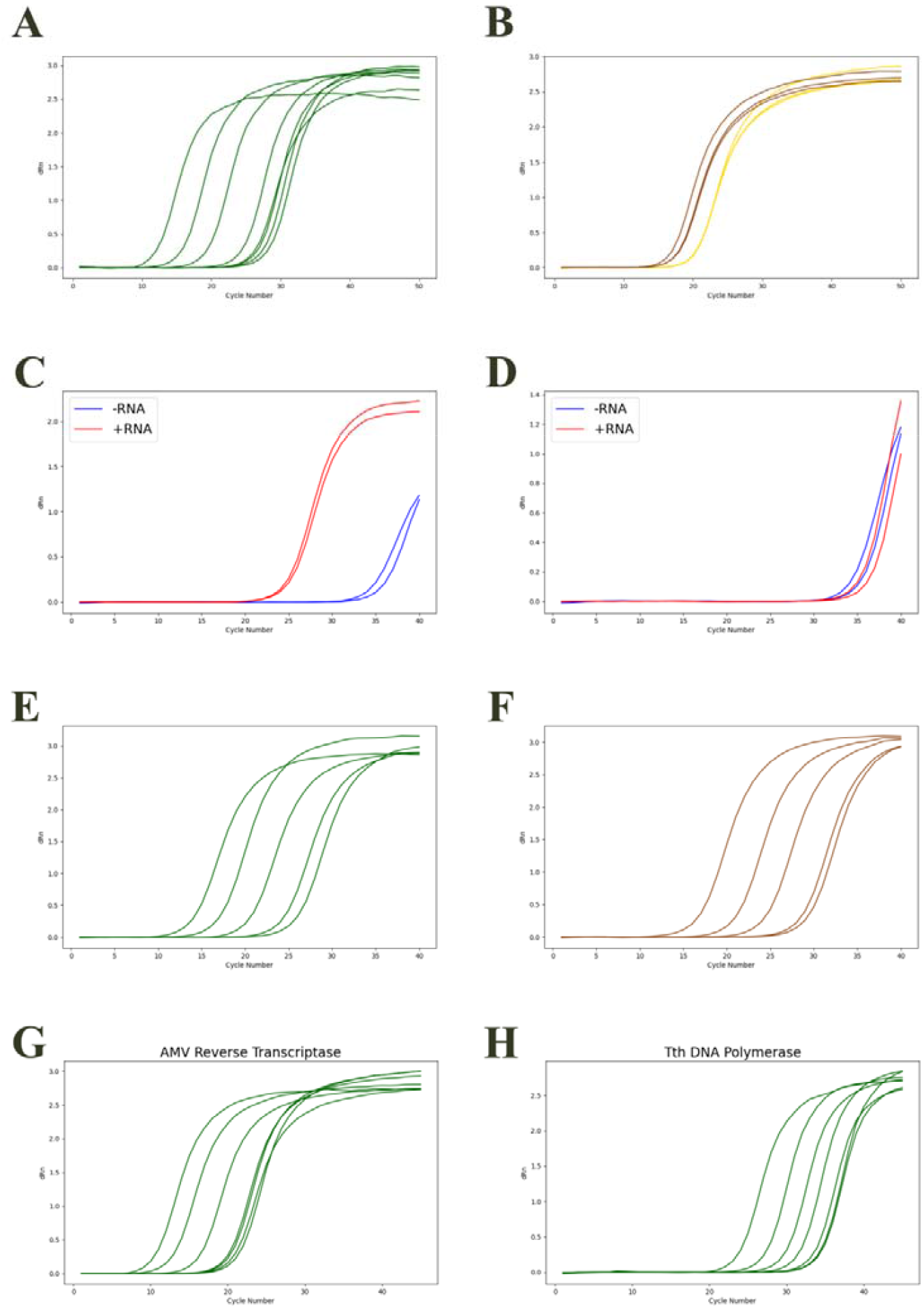
Standard Amplification Plots of CD63-Aptamer During Optimization Procedures. (A) Amplification plot with phosphorylated mediated RNA. (B) No-aptamer RT of two random RNA sequences resulted in amplification curves. Each color represents a different sequence. (C) Mediated RNA was amplified in qPCR as a template. (D) After RNase degradation, mediated RNA amplification in qPCR was eliminated. (E, F) Comparison of qPCR amplification without (E) and with (F) an RNase degradation step after RT. (G) Amplification plot of AMV Reverse Transcriptase-based RT-qPCR. (H) Amplification plot of Tth DNA Polymerase-based RT-qPCR.

#### Mediated RNA Directly Amplificated in qPCR

The potential for mediated RNA to be amplified in qPCR was investigated. When CD63-aptamer mediated RNA was directly added as a template in qPCR, an amplification curve was observed (Figure 5C). In contrast, when the mediated RNA was treated with RNase A, no significant amplification occurred (Figure 5D), indicating that RNA degradation prevented amplification.

To minimize the potential influence of mediated RNA, we included an RNase A degradation step. As shown in Figure 5E and Figure 5F, compared to reactions without RNase treatment, the amplification curves with RNase A became more orderly and shifted to the right, indicating that the mediated RNA of the CD63 aptamer interfered with qPCR. Interestingly, the amplification curves of the CD9 aptamer were largely unaffected by RNase A (data not shown), possibly because its mediated RNA contained only a single primer site and thus the cDNA of mediated RNA could not be amplificated in qPCR.

#### No-Primer Reverse Transcription also Occurred with AMV Reverse Transcriptase and Tth DNA Polymerase

To figure out whether the no-primer RT is specific to MMLV Reverse Transcriptase (RTase), AMV RTase and Tth DNA Polymerase were also tested (Figure 5G and 5H). The amplification curves for both enzymes were similar to those of MMLV RTase. The rightward shift of Tth’s curves indicated that its reverse transcription activity was relatively lower than that of MMLV RTase and AMV RTase. All three enzymes were capable of reverse transcription in the absence of a primer.

#### RNase R Enhanced the Detection Sensitivity of CD63-Aptamer Quantification

RNase R is a 3’→5’ exoribonuclease that specifically degrades single-stranded RNA.^19^ Following RNase R treatment, unbound mediated RNA is selectively digested, while mediated RNA stably bound to the aptamer remains intact. This selective degradation effectively reduces background noise and enhances signal specificity. As a result, the application of RNase R improved the detection precision by an order of magnitude (Figure 6).

**Figure 6.**
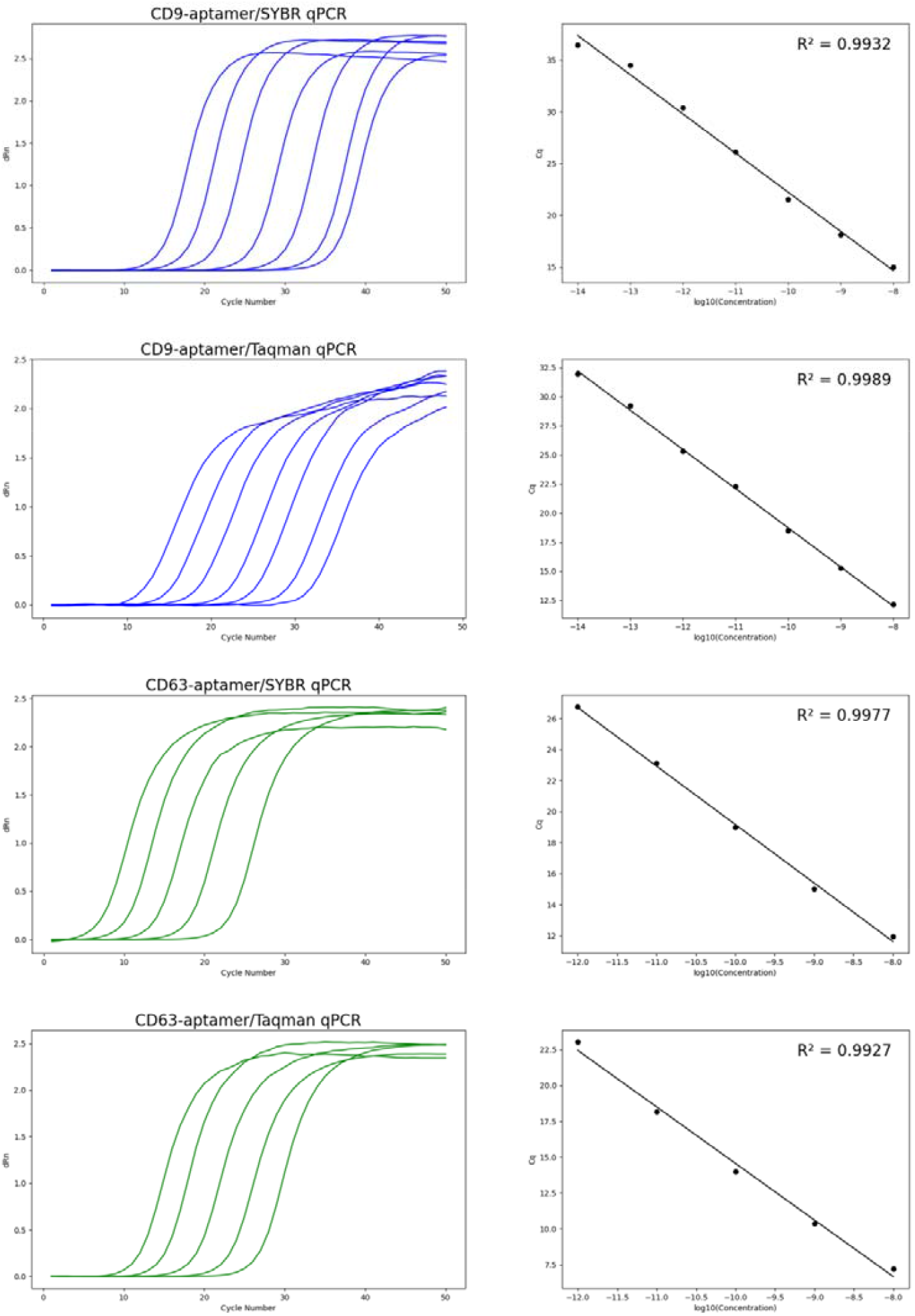
Amplification Plots and Standard Curves of RT-qPCR-Based Quantification of CD9-Aptamer (10^-8^ ∼ 10^-14^ M) and CD63-Aptamer (10^-8^ ∼ 10^-12^ M).

#### CD9-Aptamer and CD63-Aptamer Were Quantified Using SYBR Green and TaqMan qPCR

Finally, aptamer quantification methods were established. Both the SYBR Green and TaqMan methods were tested (Figure 6), and the standard curves demonstrated good linearity. For the CD9 aptamer, the limit of detection (LOD) reached 10^-14^ M, while for the CD63 aptamer, the LOD reached 10^-11^ M.

#### Quantification of CD9-Aptamer by One-step RT-qPCR

Additionally, one-step RT-qPCR was evaluated and demonstrated comparable performance to the two-step RT-qPCR method (Figure 7). Sanchita Bhadra et al.^20^ reported that Taq polymerase exhibits RTase activity, enabling one-step RT-qPCR using a single enzyme in the presence of a specific buffer. We tested two Taq polymerases from different manufacturers and confirmed their RT-qPCR capability; however, reaction conditions still required optimization (Supplementary 1 Figure 1).

**Figure 7.**
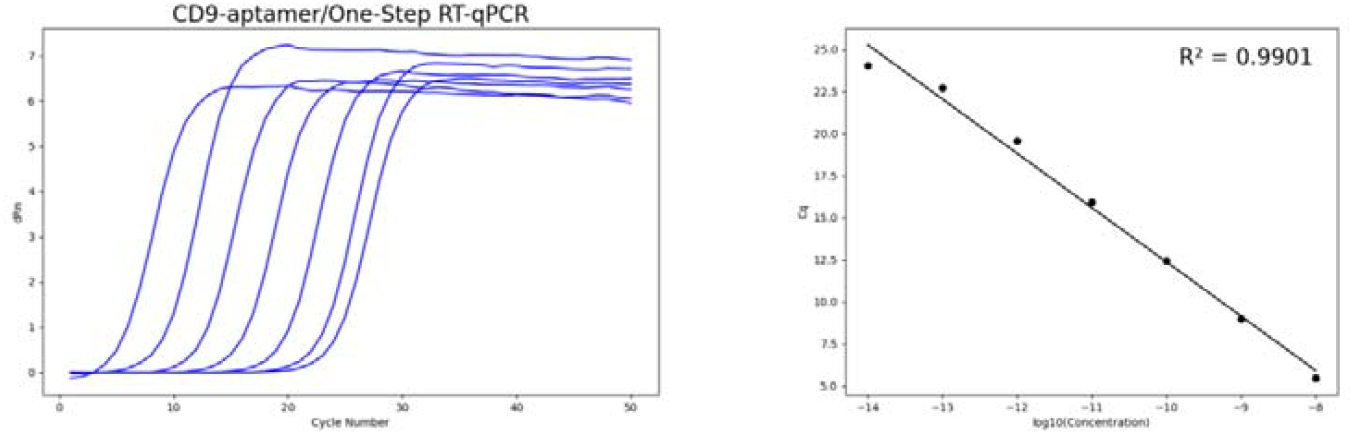
Amplification Plots and Standard Curves of One-Step RT-qPCR Quantification of CD9-Aptamer (10^-8^ ∼ 10^-14^ M).

## Discussion

In this study, we developed a novel method to quantify aptamers using RT-qPCR. Our approach does not require any structural modifications to the aptamer prior to quantification, which is a significant advantage over previous techniques.

To facilitate primer and probe design, we developed a software tool based on standardized primer design criteria. However, existing criteria from the literature were often vague or inconsistent. Therefore, we refined and explicitly defined these criteria within our software. For instance, a common recommendation is that the last bases of primer pairs should not be complementary, yet definitions of “last bases” and “complementary” are frequently unclear. In our tool, this rule is clearly defined such that the last x bases of primer pairs must not contain complementary sequences exceeding y bases. Although the default parameters performed effectively in our tests, they are flexible and can be adjusted to meet specific experimental requirements.

During the reverse transcription step, we observed an interesting phenomenon: RNA was found to serve as a substrate in the qPCR reaction (Figure 5C). Previous studies have reported that Taq polymerase exhibits reverse transcriptase activity.^20^ To mitigate the potential influence of mediated RNA, we introduced RNase A to degrade the RNA.

The most puzzling aspect of our method development was the discrepancy in qPCR results between the two aptamers tested (Figure 4). We hypothesized that this difference was due to the sequence variations between the aptamers. The mediated RNA of the CD9 aptamer had only one primer site, with the other located within the aptamer sequence. Without the aptamer, the reverse transcription product would have only one primer and could not be amplified in qPCR. In contrast, the mediated RNA of the CD63 aptamer contained two primer sites, allowing the reverse transcription product of the mediated RNA to be directly amplified in qPCR. It appeared that both mediated RNAs generated products during reverse transcription; however, only the mediated RNA of the CD63 aptamer could be amplified in qPCR without the presence of the aptamer.

The amplification curve for the reverse transcription product without the aptamer in qPCR confirmed that reverse transcription occurred even in the absence of a primer. It has been previously reported that RNA can self-prime in reverse transcription.^21,22^ To prevent self-priming, we introduced several modifications, including phosphorylation, C18, and poly-U at the 3’ end of the RNA. Although the result improved, it still showed that even in the absence of a 3’-OH group on the RNA sequence, reverse transcription occurred without the presence of the aptamer.

The only other potential source of 3’-OH in the reaction was the dNTPs. It has been reported that MMLV RTase adds a tail to DNA ends^23^, which led us to speculate that it could also add dNTPs onto other dNTPs, forming random sequences that might act as primers. However, this hypothesis was preliminarily ruled out. Mass spectrometry analysis of a 30-minute reverse transcription reaction without primers or templates did not detect any oligonucleotides (data not shown), suggesting that reverse transcription could still occur without primers.

We also tested several additives, such as DMSO, EDTA, and low molecular weight heparin, to adjust the binding strength of the double helix and reduce non-specific binding. The goal was to find an optimal concentration that would inhibit no-primer reverse transcription without interfering with aptamer-primed reactions. Unfortunately, these additives either muddied the results or inhibited reactions equally at both high and low aptamer concentrations (data not shown). Additionally, testing other enzymes, such as MMLV RTase (RNase H-, data not shown) AMV RTase and Tth DNA Polymerase, yielded similar results as MMLV RTase.

Another strategy was to degrade unbound mediated RNA, ensuring that only mediated RNA bound to the aptamer would undergo reverse transcription. RNase R treatment improved accuracy; however, it could not completely eliminate all unbound mediated RNA. We hypothesize that a subset of mediated RNA molecules may form secondary structures that lack accessible 3’ tails, which are essential for RNase R binding and degradation.

Based on these observations, we conclude that further improvement of this method requires the identification or development of a reverse transcriptase that does not catalyze reverse transcription in the absence of a primer, even at high RNA concentrations. Matej Zabrady et al.^24^ reported that HIV RTase can initiate reverse transcription without a primer. Given their structural similarity^25^, it is plausible that MMLV RTase can also initiate primer-independent reverse transcription. In their study, the primer-free reaction required initiation with a dGTP. This suggests that, in theory, it may be possible to engineer RTase variants that eliminate G-initiated, primer-independent activity without affecting primer-dependent reverse transcription, as the G-start mechanism is not necessary for regular primer-initiated RT.

This quantification method represents a significant advancement over existing techniques. For example, the LOD for CD63 aptamer quantification using our method was approximately 1 pM, and for CD9 aptamer was 0.01 pM, whereas in a pharmacokinetics study, the LOD for a different aptamer using UPLC was about 0.14 uM.^26^

Compared to the no-primer-site type (e.g., CD63 aptamer), the one-primer-site type (e.g., CD9 aptamer) achieved a significantly lower limit of detection (LOD). Therefore, we recommend attempting the one-primer-site method, even if it requires slightly relaxing restrictions in the software design process.

During re-validation of the method, we identified two key factors that may affect repeatability. First, the mediated RNA was relatively prone to degradation, so it should be divided into small volumes and replaced frequently. Second, at low concentrations, aptamers tend to adhere to tube walls, leading to sample loss during the experiment. To prevent this, carrier nucleic acids can be added to stabilize aptamers. For instance, salmon sperm DNA is commonly used to stabilize miRNA^27^. In this study, we found that other aptamers could serve the same role, such as using CD63 aptamer to protect CD9 aptamer.

Beyond the experimental details, our newly developed aptamer quantification method is precise, universal, and cost-effective, addressing a significant gap in aptamer quantification. This advancement supports broader and more effective applications of aptamers across various fields.

For example, aptamers have great potential as next-generation therapeutics, where accurate pharmacokinetic quantification is essential for clinical development. Existing methods, such as HPLC and radio-labeling, are costly and relatively inaccurate. Our RT-qPCR-based approach significantly improves precision, facilitating progress in precision medicine.

In addition to therapeutics, aptamers have promising applications in diagnostics, such as ELASA, where aptamers replace antibodies for protein detection. Traditionally, ELASA uses enzyme-based signal amplification (e.g., HRP), similar to ELISA. Our RT-qPCR quantification method could replace these enzyme-based amplification strategies, potentially improving sensitivity and accuracy in diagnostic applications.

