## Supplementary 1 for "A Universal RT-qPCR Method for DNA Aptamer Quantification"

Sup Table 1 Primer pairs selected by "Apt Primer Finder". Yellow means no primer dimer in qPCR.

| Name | Sequence |
| --- | --- |
| CD9-Apt1-F0822-1 | ATAGTCCCTTGGCGTGCTT |
| CD9-Apt1-R0822-1a | CTGTCGTCGTTTGTCTGTGTTG |
| CD9-Apt1-R0822-1b | ACCTTTCGGCTTCTCGTGTT |
| CD9-Apt1-R0822-1c | CATACCCGTCCGTCTTCTTGTT |
| CD9-Apt1-F0822-2 | CCTTGGCGTGCTTCACA |
| CD9-Apt1-R0822-2a | AACTCCCTCGGCTAACTTCT |
| CD9-Apt1-R0822-2b | CTCCTTTCCTCCCTTCAGACTT |
| CD9-Apt1-R0822-2c | AACCCTAACTTCGGCTCCTT |

Sup Table 2 Primer pairs selected by "Aptamer Primer Pair Filter". Yellow means no primer dimer in qPCR.

| Name | Sequence |
| --- | --- |
| CD63-Apt1-F0822-1 | CTGCCTGGTATGTTGCTTCTTG |
| CD63-Apt1-R0822-1 | TTCCTCGCTTTCTCGCTCTT |
| CD63-Apt1-F0822-2 | TGCTGTTGTATCGGCTCCTT |
| CD63-Apt1-R0822-2 | TGACCATCCATCTTCCCTCTTG |
| CD63-Apt1-F0822-3 | ACCTCATCATCGCCCATCAA |
| CD63-Apt1-R0822-3 | CCGTATCGTATCGCAGTAACCA |

Sup Table 3 Sequences of test RNAs and primers. “-P” means phosphorylation.

| Name | Sequence |
| --- | --- |
| Test1F | TCGCTTCTCTCGCTGATTGT |
| Test1R | TCTATGGATGCCCGTTTGGT |
| Test1RNA | UCGCUUCUCUCGCUGAUUGUACCAAACGGGCAUCCAUAGA-P |
| Test2F | TTCGTTGCCTTTCCGTTGTG |
| Test2R | AATCGCCCTGCTGTAATCCT |
| Test2RNA | UUCGUUGCCUUUCCGUUGUGAGGAUUACAGCAGGGCGAUU-P |

Sup Figure 1 Amplification Plots and Standard Curves of One-Enzyme One-Step RT-qPCR Quantification of CD9-Aptamer (10^-8^ ~ 10^-14^ M). Taq polymerase (D7211, Beyotime; M0267,NEB)
