## Supplementary 2 for "A Universal RT-qPCR Method for DNA Aptamer Quantification": METADATA

Metadata-Version: 2.1
Name: attrs
Version: 23.2.0
Summary: Classes Without Boilerplate
Project-URL: Documentation, https://www.attrs.org/
Project-URL: Changelog, https://www.attrs.org/en/stable/changelog.html
Project-URL: GitHub, https://github.com/python-attrs/attrs
Project-URL: Funding, https://github.com/sponsors/hynek
Project-URL: Tidelift, https://tidelift.com/subscription/pkg/pypi-attrs?utm_source=pypi-attrs&utm_medium=pypi
License-Expression: MIT
License-File: LICENSE
Keywords: attribute,boilerplate,class
Classifier: Development Status :: 5 - Production/Stable
Classifier: License :: OSI Approved :: MIT License
Classifier: Programming Language :: Python :: 3.7
Classifier: Programming Language :: Python :: 3.8
Classifier: Programming Language :: Python :: 3.9
Classifier: Programming Language :: Python :: 3.10
Classifier: Programming Language :: Python :: 3.11
Classifier: Programming Language :: Python :: 3.12
Classifier: Programming Language :: Python :: Implementation :: CPython
Classifier: Programming Language :: Python :: Implementation :: PyPy
Classifier: Typing :: Typed
Requires-Python: >=3.7
Requires-Dist: importlib-metadata; python_version < '3.8'
Provides-Extra: cov
Requires-Dist: attrs[tests]; extra == 'cov'
Requires-Dist: coverage[toml]>=5.3; extra == 'cov'
Provides-Extra: dev
Requires-Dist: attrs[tests]; extra == 'dev'
Requires-Dist: pre-commit; extra == 'dev'
Provides-Extra: docs
Requires-Dist: furo; extra == 'docs'
Requires-Dist: myst-parser; extra == 'docs'
Requires-Dist: sphinx; extra == 'docs'
Requires-Dist: sphinx-notfound-page; extra == 'docs'
Requires-Dist: sphinxcontrib-towncrier; extra == 'docs'
Requires-Dist: towncrier; extra == 'docs'
Requires-Dist: zope-interface; extra == 'docs'
Provides-Extra: tests
Requires-Dist: attrs[tests-no-zope]; extra == 'tests'
Requires-Dist: zope-interface; extra == 'tests'
Provides-Extra: tests-mypy
Requires-Dist: mypy>=1.6; (platform_python_implementation == 'CPython' and python_version >= '3.8') and extra == 'tests-mypy'
Requires-Dist: pytest-mypy-plugins; (platform_python_implementation == 'CPython' and python_version >= '3.8') and extra == 'tests-mypy'
Provides-Extra: tests-no-zope
Requires-Dist: attrs[tests-mypy]; extra == 'tests-no-zope'
Requires-Dist: cloudpickle; (platform_python_implementation == 'CPython') and extra == 'tests-no-zope'
Requires-Dist: hypothesis; extra == 'tests-no-zope'
Requires-Dist: pympler; extra == 'tests-no-zope'
Requires-Dist: pytest-xdist[psutil]; extra == 'tests-no-zope'
Requires-Dist: pytest>=4.3.0; extra == 'tests-no-zope'
Description-Content-Type: text/markdown

<p align="center">
  <a href="https://www.attrs.org/">
    <img src="https://raw.githubusercontent.com/python-attrs/attrs/main/docs/_static/attrs_logo.svg" width="35%" alt="attrs" />
  </a>
</p>


*attrs* is the Python package that will bring back the **joy** of **writing classes** by relieving you from the drudgery of implementing object protocols (aka [dunder methods](https://www.attrs.org/en/latest/glossary.html#term-dunder-methods)).
[Trusted by NASA](https://docs.github.com/en/account-and-profile/setting-up-and-managing-your-github-profile/customizing-your-profile/personalizing-your-profile#list-of-qualifying-repositories-for-mars-2020-helicopter-contributor-achievement) for Mars missions since 2020!

Its main goal is to help you to write **concise** and **correct** software without slowing down your code.


## Sponsors

*attrs* would not be possible without our [amazing sponsors](https://github.com/sponsors/hynek).
Especially those generously supporting us at the *The Organization* tier and higher:

<p align="center">
   <a href="https://www.variomedia.de/"><img src="https://www.attrs.org/en/23.2.0/_static/sponsors/Variomedia.svg" width="200" height="60" /></a>
   <a href="https://tidelift.com/subscription/pkg/pypi-attrs?utm_source=pypi-attrs&utm_medium=referral&utm_campaign=enterprise&utm_term=repo"><img src="https://www.attrs.org/en/23.2.0/_static/sponsors/Tidelift.svg" width="200" height="60" /></a>
   <a href="https://filepreviews.io/"><img src="https://www.attrs.org/en/23.2.0/_static/sponsors/FilePreviews.svg" width="200" height="60"/></a>
</p>

<p align="center">
   <strong>Please consider <a href="https://github.com/sponsors/hynek">joining them</a> to help make <em>attrs</em>’s maintenance more sustainable!</strong>
</p>

<!-- teaser-end -->

## Example

*attrs* gives you a class decorator and a way to declaratively define the attributes on that class:

<!-- code-begin -->

```pycon
>>> from attrs import asdict, define, make_class, Factory

>>> @define
... class SomeClass:
...     a_number: int = 42
...     list_of_numbers: list[int] = Factory(list)
...
...     def hard_math(self, another_number):
...         return self.a_number + sum(self.list_of_numbers) * another_number


>>> sc = SomeClass(1, [1, 2, 3])
>>> sc
SomeClass(a_number=1, list_of_numbers=[1, 2, 3])

>>> sc.hard_math(3)
19
>>> sc == SomeClass(1, [1, 2, 3])
True
>>> sc != SomeClass(2, [3, 2, 1])
True

>>> asdict(sc)
{'a_number': 1, 'list_of_numbers': [1, 2, 3]}

>>> SomeClass()
SomeClass(a_number=42, list_of_numbers=[])

>>> C = make_class("C", ["a", "b"])
>>> C("foo", "bar")
C(a='foo', b='bar')
```

After *declaring* your attributes, *attrs* gives you:

- a concise and explicit overview of the class's attributes,
- a nice human-readable `__repr__`,
- equality-checking methods,
- an initializer,
- and much more,

*without* writing dull boilerplate code again and again and *without* runtime performance penalties.

**Hate type annotations**!?
No problem!
Types are entirely **optional** with *attrs*.
Simply assign `attrs.field()` to the attributes instead of annotating them with types.

---

This example uses *attrs*'s modern APIs that have been introduced in version 20.1.0, and the *attrs* package import name that has been added in version 21.3.0.
The classic APIs (`@attr.s`, `attr.ib`, plus their serious-business aliases) and the `attr` package import name will remain **indefinitely**.

Please check out [*On The Core API Names*](https://www.attrs.org/en/latest/names.html) for a more in-depth explanation.


## Data Classes

On the tin, *attrs* might remind you of `dataclasses` (and indeed, `dataclasses` [are a descendant](https://hynek.me/articles/import-attrs/) of *attrs*).
In practice it does a lot more and is more flexible.
For instance it allows you to define [special handling of NumPy arrays for equality checks](https://www.attrs.org/en/stable/comparison.html#customization), allows more ways to [plug into the initialization process](https://www.attrs.org/en/stable/init.html#hooking-yourself-into-initialization), and allows for stepping through the generated methods using a debugger.

For more details, please refer to our [comparison page](https://www.attrs.org/en/stable/why.html#data-classes).


## Project Information

- [**Changelog**](https://www.attrs.org/en/stable/changelog.html)
- [**Documentation**](https://www.attrs.org/)
- [**PyPI**](https://pypi.org/project/attrs/)
- [**Source Code**](https://github.com/python-attrs/attrs)
- [**Contributing**](https://github.com/python-attrs/attrs/blob/main/.github/CONTRIBUTING.md)
- [**Third-party Extensions**](https://github.com/python-attrs/attrs/wiki/Extensions-to-attrs)
- **Get Help**: please use the `python-attrs` tag on [StackOverflow](https://stackoverflow.com/questions/tagged/python-attrs)


### *attrs* for Enterprise

Available as part of the Tidelift Subscription.

The maintainers of *attrs* and thousands of other packages are working with Tidelift to deliver commercial support and maintenance for the open source packages you use to build your applications.
Save time, reduce risk, and improve code health, while paying the maintainers of the exact packages you use.
[Learn more.](https://tidelift.com/subscription/pkg/pypi-attrs?utm_source=pypi-attrs&utm_medium=referral&utm_campaign=enterprise&utm_term=repo)

## Release Information

### Changes

- The type annotation for `attrs.resolve_types()` is now correct.
  [#1141](https://github.com/python-attrs/attrs/issues/1141)
- Type stubs now use `typing.dataclass_transform` to decorate dataclass-like decorators, instead of the non-standard `__dataclass_transform__` special form, which is only supported by Pyright.
  [#1158](https://github.com/python-attrs/attrs/issues/1158)
- Fixed serialization of namedtuple fields using `attrs.asdict/astuple()` with `retain_collection_types=True`.
  [#1165](https://github.com/python-attrs/attrs/issues/1165)
- `attrs.AttrsInstance` is now a `typing.Protocol` in both type hints and code.
  This allows you to subclass it along with another `Protocol`.
  [#1172](https://github.com/python-attrs/attrs/issues/1172)
- If *attrs* detects that `__attrs_pre_init__` accepts more than just `self`, it will call it with the same arguments as `__init__` was called.
  This allows you to, for example, pass arguments to `super().__init__()`.
  [#1187](https://github.com/python-attrs/attrs/issues/1187)
- Slotted classes now transform `functools.cached_property` decorated methods to support equivalent semantics.
  [#1200](https://github.com/python-attrs/attrs/issues/1200)
- Added *class_body* argument to `attrs.make_class()` to provide additional attributes for newly created classes.
  It is, for example, now possible to attach methods.
  [#1203](https://github.com/python-attrs/attrs/issues/1203)


---

[Full changelog](https://www.attrs.org/en/stable/changelog.html)
