## Supplementary 2 for "A Universal RT-qPCR Method for DNA Aptamer Quantification": README_STARTUP

This is the IPython startup directory

.py and .ipy files in this directory will be run *prior* to any code or files specified
via the exec_lines or exec_files configurables whenever you load this profile.

Files will be run in lexicographical order, so you can control the execution order of files
with a prefix, e.g.::

    00-first.py
    50-middle.py
    99-last.ipy
