## Supplementary 2 for "A Universal RT-qPCR Method for DNA Aptamer Quantification": autoreload.py

```
"""IPython extension to reload modules before executing user code.

``autoreload`` reloads modules automatically before entering the execution of
code typed at the IPython prompt.

This makes for example the following workflow possible:

.. sourcecode:: ipython

   In [1]: %load_ext autoreload

   In [2]: %autoreload 2

   In [3]: from foo import some_function

   In [4]: some_function()
   Out[4]: 42

   In [5]: # open foo.py in an editor and change some_function to return 43

   In [6]: some_function()
   Out[6]: 43

The module was reloaded without reloading it explicitly, and the object
imported with ``from foo import ...`` was also updated.

Usage
=====

The following magic commands are provided:

``%autoreload``, ``%autoreload now``

    Reload all modules (except those excluded by ``%aimport``)
    automatically now.

``%autoreload 0``, ``%autoreload off``

    Disable automatic reloading.

``%autoreload 1``, ``%autoreload explicit``

    Reload all modules imported with ``%aimport`` every time before
    executing the Python code typed.

``%autoreload 2``, ``%autoreload all``

    Reload all modules (except those excluded by ``%aimport``) every
    time before executing the Python code typed.

``%autoreload 3``, ``%autoreload complete``

    Same as 2/all, but also adds any new objects in the module. See
    unit test at IPython/extensions/tests/test_autoreload.py::test_autoload_newly_added_objects

  Adding ``--print`` or ``-p`` to the ``%autoreload`` line will print autoreload activity to
  standard out. ``--log`` or ``-l`` will do it to the log at INFO level; both can be used
  simultaneously.

``%aimport``

    List modules which are to be automatically imported or not to be imported.

``%aimport foo``

    Import module 'foo' and mark it to be autoreloaded for ``%autoreload 1``

``%aimport foo, bar``

    Import modules 'foo', 'bar' and mark them to be autoreloaded for ``%autoreload 1``

``%aimport -foo``

    Mark module 'foo' to not be autoreloaded.

Caveats
=======

Reloading Python modules in a reliable way is in general difficult,
and unexpected things may occur. ``%autoreload`` tries to work around
common pitfalls by replacing function code objects and parts of
classes previously in the module with new versions. This makes the
following things to work:

- Functions and classes imported via 'from xxx import foo' are upgraded
  to new versions when 'xxx' is reloaded.

- Methods and properties of classes are upgraded on reload, so that
  calling 'c.foo()' on an object 'c' created before the reload causes
  the new code for 'foo' to be executed.

Some of the known remaining caveats are:

- Replacing code objects does not always succeed: changing a @property
  in a class to an ordinary method or a method to a member variable
  can cause problems (but in old objects only).

- Functions that are removed (eg. via monkey-patching) from a module
  before it is reloaded are not upgraded.

- C extension modules cannot be reloaded, and so cannot be autoreloaded.

- While comparing Enum and Flag, the 'is' Identity Operator is used (even in the case '==' has been used (Similar to the 'None' keyword)).

- Reloading a module, or importing the same module by a different name, creates new Enums. These may look the same, but are not.
"""

from IPython.core import magic_arguments
from IPython.core.magic import Magics, magics_class, line_magic

__skip_doctest__ = True

# -----------------------------------------------------------------------------
#  Copyright (C) 2000 Thomas Heller
#  Copyright (C) 2008 Pauli Virtanen <>
#  Copyright (C) 2012  The IPython Development Team
#
#  Distributed under the terms of the BSD License.  The full license is in
#  the file COPYING, distributed as part of this software.
# -----------------------------------------------------------------------------
#
# This IPython module is written by Pauli Virtanen, based on the autoreload
# code by Thomas Heller.

# -----------------------------------------------------------------------------
# Imports
# -----------------------------------------------------------------------------

import os
import sys
import traceback
import types
import weakref
import gc
import logging
from importlib import import_module, reload
from importlib.util import source_from_cache

# ------------------------------------------------------------------------------
# Autoreload functionality
# ------------------------------------------------------------------------------


class ModuleReloader:
    enabled = False
    """Whether this reloader is enabled"""

    check_all = True
    """Autoreload all modules, not just those listed in 'modules'"""

    autoload_obj = False
    """Autoreload all modules AND autoload all new objects"""

    def __init__(self, shell=None):
        # Modules that failed to reload: {module: mtime-on-failed-reload, ...}
        self.failed = {}
        # Modules specially marked as autoreloadable.
        self.modules = {}
        # Modules specially marked as not autoreloadable.
        self.skip_modules = {}
        # (module-name, name) -> weakref, for replacing old code objects
        self.old_objects = {}
        # Module modification timestamps
        self.modules_mtimes = {}
        self.shell = shell

        # Reporting callable for verbosity
        self._report = lambda msg: None  # by default, be quiet.

        # Cache module modification times
        self.check(check_all=True, do_reload=False)

        # To hide autoreload errors
        self.hide_errors = False

    def mark_module_skipped(self, module_name):
        """Skip reloading the named module in the future"""
        try:
            del self.modules[module_name]
        except KeyError:
            pass
        self.skip_modules[module_name] = True

    def mark_module_reloadable(self, module_name):
        """Reload the named module in the future (if it is imported)"""
        try:
            del self.skip_modules[module_name]
        except KeyError:
            pass
        self.modules[module_name] = True

    def aimport_module(self, module_name):
        """Import a module, and mark it reloadable

        Returns
        -------
        top_module : module
            The imported module if it is top-level, or the top-level
        top_name : module
            Name of top_module

        """
        self.mark_module_reloadable(module_name)

        import_module(module_name)
        top_name = module_name.split(".")[0]
        top_module = sys.modules[top_name]
        return top_module, top_name

    def filename_and_mtime(self, module):
        if not hasattr(module, "__file__") or module.__file__ is None:
            return None, None

        if getattr(module, "__name__", None) in [None, "__mp_main__", "__main__"]:
            # we cannot reload(__main__) or reload(__mp_main__)
            return None, None

        filename = module.__file__
        path, ext = os.path.splitext(filename)

        if ext.lower() == ".py":
            py_filename = filename
        else:
            try:
                py_filename = source_from_cache(filename)
            except ValueError:
                return None, None

        try:
            pymtime = os.stat(py_filename).st_mtime
        except OSError:
            return None, None

        return py_filename, pymtime

    def check(self, check_all=False, do_reload=True):
        """Check whether some modules need to be reloaded."""

        if not self.enabled and not check_all:
            return

        if check_all or self.check_all:
            modules = list(sys.modules.keys())
        else:
            modules = list(self.modules.keys())

        for modname in modules:
            m = sys.modules.get(modname, None)

            if modname in self.skip_modules:
                continue

            py_filename, pymtime = self.filename_and_mtime(m)
            if py_filename is None:
                continue

            try:
                if pymtime <= self.modules_mtimes[modname]:
                    continue
            except KeyError:
                self.modules_mtimes[modname] = pymtime
                continue
            else:
                if self.failed.get(py_filename, None) == pymtime:
                    continue

            self.modules_mtimes[modname] = pymtime

            # If we've reached this point, we should try to reload the module
            if do_reload:
                self._report(f"Reloading '{modname}'.")
                try:
                    if self.autoload_obj:
                        superreload(m, reload, self.old_objects, self.shell)
                    else:
                        superreload(m, reload, self.old_objects)
                    if py_filename in self.failed:
                        del self.failed[py_filename]
                except:
                    if not self.hide_errors:
                        print(
                            "[autoreload of {} failed: {}]".format(
                                modname, traceback.format_exc(10)
                            ),
                            file=sys.stderr,
                        )
                    self.failed[py_filename] = pymtime


# ------------------------------------------------------------------------------
# superreload
# ------------------------------------------------------------------------------


func_attrs = [
    "__code__",
    "__defaults__",
    "__doc__",
    "__closure__",
    "__globals__",
    "__dict__",
]


def update_function(old, new):
    """Upgrade the code object of a function"""
    for name in func_attrs:
        try:
            setattr(old, name, getattr(new, name))
        except (AttributeError, TypeError):
            pass


def update_instances(old, new):
    """Use garbage collector to find all instances that refer to the old
    class definition and update their __class__ to point to the new class
    definition"""

    refs = gc.get_referrers(old)

    for ref in refs:
        if type(ref) is old:
            object.__setattr__(ref, "__class__", new)


def update_class(old, new):
    """Replace stuff in the __dict__ of a class, and upgrade
    method code objects, and add new methods, if any"""
    for key in list(old.__dict__.keys()):
        old_obj = getattr(old, key)
        try:
            new_obj = getattr(new, key)
            # explicitly checking that comparison returns True to handle
            # cases where `==` doesn't return a boolean.
            if (old_obj == new_obj) is True:
                continue
        except AttributeError:
            # obsolete attribute: remove it
            try:
                delattr(old, key)
            except (AttributeError, TypeError):
                pass
            continue
        except ValueError:
            # can't compare nested structures containing
            # numpy arrays using `==`
            pass

        if update_generic(old_obj, new_obj):
            continue

        try:
            setattr(old, key, getattr(new, key))
        except (AttributeError, TypeError):
            pass  # skip non-writable attributes

    for key in list(new.__dict__.keys()):
        if key not in list(old.__dict__.keys()):
            try:
                setattr(old, key, getattr(new, key))
            except (AttributeError, TypeError):
                pass  # skip non-writable attributes

    # update all instances of class
    update_instances(old, new)


def update_property(old, new):
    """Replace get/set/del functions of a property"""
    update_generic(old.fdel, new.fdel)
    update_generic(old.fget, new.fget)
    update_generic(old.fset, new.fset)


def isinstance2(a, b, typ):
    return isinstance(a, typ) and isinstance(b, typ)


UPDATE_RULES = [
    (lambda a, b: isinstance2(a, b, type), update_class),
    (lambda a, b: isinstance2(a, b, types.FunctionType), update_function),
    (lambda a, b: isinstance2(a, b, property), update_property),
]
UPDATE_RULES.extend(
    [
        (
            lambda a, b: isinstance2(a, b, types.MethodType),
            lambda a, b: update_function(a.__func__, b.__func__),
        ),
    ]
)


def update_generic(a, b):
    for type_check, update in UPDATE_RULES:
        if type_check(a, b):
            update(a, b)
            return True
    return False


class StrongRef:
    def __init__(self, obj):
        self.obj = obj

    def __call__(self):
        return self.obj


mod_attrs = [
    "__name__",
    "__doc__",
    "__package__",
    "__loader__",
    "__spec__",
    "__file__",
    "__cached__",
    "__builtins__",
]


def append_obj(module, d, name, obj, autoload=False):
    in_module = hasattr(obj, "__module__") and obj.__module__ == module.__name__
    if autoload:
        # check needed for module global built-ins
        if not in_module and name in mod_attrs:
            return False
    else:
        if not in_module:
            return False

    key = (module.__name__, name)
    try:
        d.setdefault(key, []).append(weakref.ref(obj))
    except TypeError:
        pass
    return True


def superreload(module, reload=reload, old_objects=None, shell=None):
    """Enhanced version of the builtin reload function.

    superreload remembers objects previously in the module, and

    - upgrades the class dictionary of every old class in the module
    - upgrades the code object of every old function and method
    - clears the module's namespace before reloading

    """
    if old_objects is None:
        old_objects = {}

    # collect old objects in the module
    for name, obj in list(module.__dict__.items()):
        if not append_obj(module, old_objects, name, obj):
            continue
        key = (module.__name__, name)
        try:
            old_objects.setdefault(key, []).append(weakref.ref(obj))
        except TypeError:
            pass

    # reload module
    try:
        # clear namespace first from old cruft
        old_dict = module.__dict__.copy()
        old_name = module.__name__
        module.__dict__.clear()
        module.__dict__["__name__"] = old_name
        module.__dict__["__loader__"] = old_dict["__loader__"]
    except (TypeError, AttributeError, KeyError):
        pass

    try:
        module = reload(module)
    except:
        # restore module dictionary on failed reload
        module.__dict__.update(old_dict)
        raise

    # iterate over all objects and update functions & classes
    for name, new_obj in list(module.__dict__.items()):
        key = (module.__name__, name)
        if key not in old_objects:
            # here 'shell' acts both as a flag and as an output var
            if (
                shell is None
                or name == "Enum"
                or not append_obj(module, old_objects, name, new_obj, True)
            ):
                continue
            shell.user_ns[name] = new_obj

        new_refs = []
        for old_ref in old_objects[key]:
            old_obj = old_ref()
            if old_obj is None:
                continue
            new_refs.append(old_ref)
            update_generic(old_obj, new_obj)

        if new_refs:
            old_objects[key] = new_refs
        else:
            del old_objects[key]

    return module


# ------------------------------------------------------------------------------
# IPython connectivity
# ------------------------------------------------------------------------------


@magics_class
class AutoreloadMagics(Magics):
    def __init__(self, *a, **kw):
        super().__init__(*a, **kw)
        self._reloader = ModuleReloader(self.shell)
        self._reloader.check_all = False
        self._reloader.autoload_obj = False
        self.loaded_modules = set(sys.modules)

    @line_magic
    @magic_arguments.magic_arguments()
    @magic_arguments.argument(
        "mode",
        type=str,
        default="now",
        nargs="?",
        help="""blank or 'now' - Reload all modules (except those excluded by %%aimport)
             automatically now.

             '0' or 'off' - Disable automatic reloading.

             '1' or 'explicit' - Reload only modules imported with %%aimport every
             time before executing the Python code typed.

             '2' or 'all' - Reload all modules (except those excluded by %%aimport)
             every time before executing the Python code typed.

             '3' or 'complete' - Same as 2/all, but also but also adds any new
             objects in the module.
             """,
    )
    @magic_arguments.argument(
        "-p",
        "--print",
        action="store_true",
        default=False,
        help="Show autoreload activity using `print` statements",
    )
    @magic_arguments.argument(
        "-l",
        "--log",
        action="store_true",
        default=False,
        help="Show autoreload activity using the logger",
    )
    @magic_arguments.argument(
        "--hide-errors",
        action="store_true",
        default=False,
        help="Hide autoreload errors",
    )
    def autoreload(self, line=""):
        r"""%autoreload => Reload modules automatically

        %autoreload or %autoreload now
        Reload all modules (except those excluded by %aimport) automatically
        now.

        %autoreload 0 or %autoreload off
        Disable automatic reloading.

        %autoreload 1 or %autoreload explicit
        Reload only modules imported with %aimport every time before executing
        the Python code typed.

        %autoreload 2 or %autoreload all
        Reload all modules (except those excluded by %aimport) every time
        before executing the Python code typed.

        %autoreload 3 or %autoreload complete
        Same as 2/all, but also but also adds any new objects in the module. See
        unit test at IPython/extensions/tests/test_autoreload.py::test_autoload_newly_added_objects

        The optional arguments --print and --log control display of autoreload activity. The default
        is to act silently; --print (or -p) will print out the names of modules that are being
        reloaded, and --log (or -l) outputs them to the log at INFO level.

        The optional argument --hide-errors hides any errors that can happen when trying to
        reload code.

        Reloading Python modules in a reliable way is in general
        difficult, and unexpected things may occur. %autoreload tries to
        work around common pitfalls by replacing function code objects and
        parts of classes previously in the module with new versions. This
        makes the following things to work:

        - Functions and classes imported via 'from xxx import foo' are upgraded
          to new versions when 'xxx' is reloaded.

        - Methods and properties of classes are upgraded on reload, so that
          calling 'c.foo()' on an object 'c' created before the reload causes
          the new code for 'foo' to be executed.

        Some of the known remaining caveats are:

        - Replacing code objects does not always succeed: changing a @property
          in a class to an ordinary method or a method to a member variable
          can cause problems (but in old objects only).

        - Functions that are removed (eg. via monkey-patching) from a module
          before it is reloaded are not upgraded.

        - C extension modules cannot be reloaded, and so cannot be
          autoreloaded.

        """
        args = magic_arguments.parse_argstring(self.autoreload, line)
        mode = args.mode.lower()

        p = print

        logger = logging.getLogger("autoreload")

        l = logger.info

        def pl(msg):
            p(msg)
            l(msg)

        if args.print is False and args.log is False:
            self._reloader._report = lambda msg: None
        elif args.print is True:
            if args.log is True:
                self._reloader._report = pl
            else:
                self._reloader._report = p
        elif args.log is True:
            self._reloader._report = l

        self._reloader.hide_errors = args.hide_errors

        if mode == "" or mode == "now":
            self._reloader.check(True)
        elif mode == "0" or mode == "off":
            self._reloader.enabled = False
        elif mode == "1" or mode == "explicit":
            self._reloader.enabled = True
            self._reloader.check_all = False
            self._reloader.autoload_obj = False
        elif mode == "2" or mode == "all":
            self._reloader.enabled = True
            self._reloader.check_all = True
            self._reloader.autoload_obj = False
        elif mode == "3" or mode == "complete":
            self._reloader.enabled = True
            self._reloader.check_all = True
            self._reloader.autoload_obj = True
        else:
            raise ValueError(f'Unrecognized autoreload mode "{mode}".')

    @line_magic
    def aimport(self, parameter_s="", stream=None):
        """%aimport => Import modules for automatic reloading.

        %aimport
        List modules to automatically import and not to import.

        %aimport foo
        Import module 'foo' and mark it to be autoreloaded for %autoreload explicit

        %aimport foo, bar
        Import modules 'foo', 'bar' and mark them to be autoreloaded for %autoreload explicit

        %aimport -foo, bar
        Mark module 'foo' to not be autoreloaded for %autoreload explicit, all, or complete, and 'bar'
        to be autoreloaded for mode explicit.
        """
        modname = parameter_s
        if not modname:
            to_reload = sorted(self._reloader.modules.keys())
            to_skip = sorted(self._reloader.skip_modules.keys())
            if stream is None:
                stream = sys.stdout
            if self._reloader.check_all:
                stream.write("Modules to reload:\nall-except-skipped\n")
            else:
                stream.write("Modules to reload:\n%s\n" % " ".join(to_reload))
            stream.write("\nModules to skip:\n%s\n" % " ".join(to_skip))
        else:
            for _module in [_.strip() for _ in modname.split(",")]:
                if _module.startswith("-"):
                    _module = _module[1:].strip()
                    self._reloader.mark_module_skipped(_module)
                else:
                    top_module, top_name = self._reloader.aimport_module(_module)

                    # Inject module to user namespace
                    self.shell.push({top_name: top_module})

    def pre_run_cell(self, info):
        if self._reloader.enabled:
            try:
                self._reloader.check()
            except:
                pass

    def post_execute_hook(self):
        """Cache the modification times of any modules imported in this execution"""
        newly_loaded_modules = set(sys.modules) - self.loaded_modules
        for modname in newly_loaded_modules:
            _, pymtime = self._reloader.filename_and_mtime(sys.modules[modname])
            if pymtime is not None:
                self._reloader.modules_mtimes[modname] = pymtime

        self.loaded_modules.update(newly_loaded_modules)


def load_ipython_extension(ip):
    """Load the extension in IPython."""
    auto_reload = AutoreloadMagics(ip)
    ip.register_magics(auto_reload)
    ip.events.register("pre_run_cell", auto_reload.pre_run_cell)
    ip.events.register("post_execute", auto_reload.post_execute_hook)
```
