## Supplementary 2 for "A Universal RT-qPCR Method for DNA Aptamer Quantification": storemagic.py

```
# -*- coding: utf-8 -*-
"""
%store magic for lightweight persistence.

Stores variables, aliases and macros in IPython's database.

To automatically restore stored variables at startup, add this to your
:file:`ipython_config.py` file::

  c.StoreMagics.autorestore = True
"""

# Copyright (c) IPython Development Team.
# Distributed under the terms of the Modified BSD License.

import inspect, os, sys, textwrap

from IPython.core.error import UsageError
from IPython.core.magic import Magics, magics_class, line_magic
from IPython.testing.skipdoctest import skip_doctest
from traitlets import Bool


def restore_aliases(ip, alias=None):
    staliases = ip.db.get('stored_aliases', {})
    if alias is None:
        for k,v in staliases.items():
            #print "restore alias",k,v # dbg
            #self.alias_table[k] = v
            ip.alias_manager.define_alias(k,v)
    else:
        ip.alias_manager.define_alias(alias, staliases[alias])


def refresh_variables(ip):
    db = ip.db
    for key in db.keys('autorestore/*'):
        # strip autorestore
        justkey = os.path.basename(key)
        try:
            obj = db[key]
        except KeyError:
            print("Unable to restore variable '%s', ignoring (use %%store -d to forget!)" % justkey)
            print("The error was:", sys.exc_info()[0])
        else:
            #print "restored",justkey,"=",obj #dbg
            ip.user_ns[justkey] = obj


def restore_dhist(ip):
    ip.user_ns['_dh'] = ip.db.get('dhist',[])


def restore_data(ip):
    refresh_variables(ip)
    restore_aliases(ip)
    restore_dhist(ip)


@magics_class
class StoreMagics(Magics):
    """Lightweight persistence for python variables.

    Provides the %store magic."""

    autorestore = Bool(False, help=
        """If True, any %store-d variables will be automatically restored
        when IPython starts.
        """
    ).tag(config=True)

    def __init__(self, shell):
        super(StoreMagics, self).__init__(shell=shell)
        self.shell.configurables.append(self)
        if self.autorestore:
            restore_data(self.shell)

    @skip_doctest
    @line_magic
    def store(self, parameter_s=''):
        """Lightweight persistence for python variables.

        Example::

          In [1]: l = ['hello',10,'world']
          In [2]: %store l
          Stored 'l' (list)
          In [3]: exit

          (IPython session is closed and started again...)

          ville@badger:~$ ipython
          In [1]: l
          NameError: name 'l' is not defined
          In [2]: %store -r
          In [3]: l
          Out[3]: ['hello', 10, 'world']

        Usage:

        * ``%store``          - Show list of all variables and their current
                                values
        * ``%store spam bar`` - Store the *current* value of the variables spam
                                and bar to disk
        * ``%store -d spam``  - Remove the variable and its value from storage
        * ``%store -z``       - Remove all variables from storage
        * ``%store -r``       - Refresh all variables, aliases and directory history
                                from store (overwrite current vals)
        * ``%store -r spam bar`` - Refresh specified variables and aliases from store
                                   (delete current val)
        * ``%store foo >a.txt``  - Store value of foo to new file a.txt
        * ``%store foo >>a.txt`` - Append value of foo to file a.txt

        It should be noted that if you change the value of a variable, you
        need to %store it again if you want to persist the new value.

        Note also that the variables will need to be pickleable; most basic
        python types can be safely %store'd.

        Also aliases can be %store'd across sessions.
        To remove an alias from the storage, use the %unalias magic.
        """

        opts,argsl = self.parse_options(parameter_s,'drz',mode='string')
        args = argsl.split()
        ip = self.shell
        db = ip.db
        # delete
        if 'd' in opts:
            try:
                todel = args[0]
            except IndexError as e:
                raise UsageError('You must provide the variable to forget') from e
            else:
                try:
                    del db['autorestore/' + todel]
                except BaseException as e:
                    raise UsageError("Can't delete variable '%s'" % todel) from e
        # reset
        elif 'z' in opts:
            for k in db.keys('autorestore/*'):
                del db[k]

        elif 'r' in opts:
            if args:
                for arg in args:
                    try:
                        obj = db["autorestore/" + arg]
                    except KeyError:
                        try:
                            restore_aliases(ip, alias=arg)
                        except KeyError:
                            print("no stored variable or alias %s" % arg)
                    else:
                        ip.user_ns[arg] = obj
            else:
                restore_data(ip)

        # run without arguments -> list variables & values
        elif not args:
            vars = db.keys('autorestore/*')
            vars.sort()
            if vars:
                size = max(map(len, vars))
            else:
                size = 0

            print('Stored variables and their in-db values:')
            fmt = '%-'+str(size)+'s -> %s'
            get = db.get
            for var in vars:
                justkey = os.path.basename(var)
                # print 30 first characters from every var
                print(fmt % (justkey, repr(get(var, '<unavailable>'))[:50]))

        # default action - store the variable
        else:
            # %store foo >file.txt or >>file.txt
            if len(args) > 1 and args[1].startswith(">"):
                fnam = os.path.expanduser(args[1].lstrip(">").lstrip())
                if args[1].startswith(">>"):
                    fil = open(fnam, "a", encoding="utf-8")
                else:
                    fil = open(fnam, "w", encoding="utf-8")
                with fil:
                    obj = ip.ev(args[0])
                    print("Writing '%s' (%s) to file '%s'." % (args[0],
                        obj.__class__.__name__, fnam))

                    if not isinstance (obj, str):
                        from pprint import pprint
                        pprint(obj, fil)
                    else:
                        fil.write(obj)
                        if not obj.endswith('\n'):
                            fil.write('\n')

                return

            # %store foo
            for arg in args:
                try:
                    obj = ip.user_ns[arg]
                except KeyError:
                    # it might be an alias
                    name = arg
                    try:
                        cmd = ip.alias_manager.retrieve_alias(name)
                    except ValueError as e:
                        raise UsageError("Unknown variable '%s'" % name) from e

                    staliases = db.get('stored_aliases',{})
                    staliases[name] = cmd
                    db['stored_aliases'] = staliases
                    print("Alias stored: %s (%s)" % (name, cmd))
                    return

                else:
                    modname = getattr(inspect.getmodule(obj), '__name__', '')
                    if modname == '__main__':
                        print(textwrap.dedent("""\
                        Warning:%s is %s
                        Proper storage of interactively declared classes (or instances
                        of those classes) is not possible! Only instances
                        of classes in real modules on file system can be %%store'd.
                        """ % (arg, obj) ))
                        return
                    #pickled = pickle.dumps(obj)
                    db[ 'autorestore/' + arg ] = obj
                    print("Stored '%s' (%s)" % (arg, obj.__class__.__name__))


def load_ipython_extension(ip):
    """Load the extension in IPython."""
    ip.register_magics(StoreMagics)
```
