## Supplementary 2 for "A Universal RT-qPCR Method for DNA Aptamer Quantification": test_autoreload.py

```
"""Tests for autoreload extension.
"""
# -----------------------------------------------------------------------------
#  Copyright (c) 2012 IPython Development Team.
#
#  Distributed under the terms of the Modified BSD License.
#
#  The full license is in the file COPYING.txt, distributed with this software.
# -----------------------------------------------------------------------------

# -----------------------------------------------------------------------------
# Imports
# -----------------------------------------------------------------------------

import os
import platform
import pytest
import sys
import tempfile
import textwrap
import shutil
import random
import time
import traceback
from io import StringIO
from dataclasses import dataclass

import IPython.testing.tools as tt

from unittest import TestCase

from IPython.extensions.autoreload import AutoreloadMagics
from IPython.core.events import EventManager, pre_run_cell
from IPython.testing.decorators import skipif_not_numpy
from IPython.core.interactiveshell import ExecutionInfo

if platform.python_implementation() == "PyPy":
    pytest.skip(
        "Current autoreload implementation is extremely slow on PyPy",
        allow_module_level=True,
    )

# -----------------------------------------------------------------------------
# Test fixture
# -----------------------------------------------------------------------------

noop = lambda *a, **kw: None


class FakeShell:
    def __init__(self):
        self.ns = {}
        self.user_ns = self.ns
        self.user_ns_hidden = {}
        self.events = EventManager(self, {"pre_run_cell", pre_run_cell})
        self.auto_magics = AutoreloadMagics(shell=self)
        self.events.register("pre_run_cell", self.auto_magics.pre_run_cell)

    register_magics = set_hook = noop

    def showtraceback(
        self,
        exc_tuple=None,
        filename=None,
        tb_offset=None,
        exception_only=False,
        running_compiled_code=False,
    ):
        traceback.print_exc()

    def run_code(self, code):
        self.events.trigger(
            "pre_run_cell",
            ExecutionInfo(
                raw_cell="",
                store_history=False,
                silent=False,
                shell_futures=False,
                cell_id=None,
            ),
        )
        exec(code, self.user_ns)
        self.auto_magics.post_execute_hook()

    def push(self, items):
        self.ns.update(items)

    def magic_autoreload(self, parameter):
        self.auto_magics.autoreload(parameter)

    def magic_aimport(self, parameter, stream=None):
        self.auto_magics.aimport(parameter, stream=stream)
        self.auto_magics.post_execute_hook()


class Fixture(TestCase):
    """Fixture for creating test module files"""

    test_dir = None
    old_sys_path = None
    filename_chars = "abcdefghijklmopqrstuvwxyz0123456789"

    def setUp(self):
        self.test_dir = tempfile.mkdtemp()
        self.old_sys_path = list(sys.path)
        sys.path.insert(0, self.test_dir)
        self.shell = FakeShell()

    def tearDown(self):
        shutil.rmtree(self.test_dir)
        sys.path = self.old_sys_path

        self.test_dir = None
        self.old_sys_path = None
        self.shell = None

    def get_module(self):
        module_name = "tmpmod_" + "".join(random.sample(self.filename_chars, 20))
        if module_name in sys.modules:
            del sys.modules[module_name]
        file_name = os.path.join(self.test_dir, module_name + ".py")
        return module_name, file_name

    def write_file(self, filename, content):
        """
        Write a file, and force a timestamp difference of at least one second

        Notes
        -----
        Python's .pyc files record the timestamp of their compilation
        with a time resolution of one second.

        Therefore, we need to force a timestamp difference between .py
        and .pyc, without having the .py file be timestamped in the
        future, and without changing the timestamp of the .pyc file
        (because that is stored in the file).  The only reliable way
        to achieve this seems to be to sleep.
        """
        content = textwrap.dedent(content)
        # Sleep one second + eps
        time.sleep(1.05)

        # Write
        with open(filename, "w", encoding="utf-8") as f:
            f.write(content)

    def new_module(self, code):
        code = textwrap.dedent(code)
        mod_name, mod_fn = self.get_module()
        with open(mod_fn, "w", encoding="utf-8") as f:
            f.write(code)
        return mod_name, mod_fn


# -----------------------------------------------------------------------------
# Test automatic reloading
# -----------------------------------------------------------------------------


def pickle_get_current_class(obj):
    """
    Original issue comes from pickle; hence the name.
    """
    name = obj.__class__.__name__
    module_name = getattr(obj, "__module__", None)
    obj2 = sys.modules[module_name]
    for subpath in name.split("."):
        obj2 = getattr(obj2, subpath)
    return obj2


class TestAutoreload(Fixture):
    def test_reload_enums(self):
        mod_name, mod_fn = self.new_module(
            textwrap.dedent(
                """
                                from enum import Enum
                                class MyEnum(Enum):
                                    A = 'A'
                                    B = 'B'
                            """
            )
        )
        self.shell.magic_autoreload("2")
        self.shell.magic_aimport(mod_name)
        self.write_file(
            mod_fn,
            textwrap.dedent(
                """
                                from enum import Enum
                                class MyEnum(Enum):
                                    A = 'A'
                                    B = 'B'
                                    C = 'C'
                            """
            ),
        )
        with tt.AssertNotPrints(
            ("[autoreload of %s failed:" % mod_name), channel="stderr"
        ):
            self.shell.run_code("pass")  # trigger another reload

    def test_reload_class_type(self):
        self.shell.magic_autoreload("2")
        mod_name, mod_fn = self.new_module(
            """
            class Test():
                def meth(self):
                    return "old"
        """
        )
        assert "test" not in self.shell.ns
        assert "result" not in self.shell.ns

        self.shell.run_code("from %s import Test" % mod_name)
        self.shell.run_code("test = Test()")

        self.write_file(
            mod_fn,
            """
            class Test():
                def meth(self):
                    return "new"
        """,
        )

        test_object = self.shell.ns["test"]

        # important to trigger autoreload logic !
        self.shell.run_code("pass")

        test_class = pickle_get_current_class(test_object)
        assert isinstance(test_object, test_class)

        # extra check.
        self.shell.run_code("import pickle")
        self.shell.run_code("p = pickle.dumps(test)")

    def test_reload_class_attributes(self):
        self.shell.magic_autoreload("2")
        mod_name, mod_fn = self.new_module(
            textwrap.dedent(
                """
                                class MyClass:

                                    def __init__(self, a=10):
                                        self.a = a
                                        self.b = 22 
                                        # self.toto = 33

                                    def square(self):
                                        print('compute square')
                                        return self.a*self.a
                            """
            )
        )
        self.shell.run_code("from %s import MyClass" % mod_name)
        self.shell.run_code("first = MyClass(5)")
        self.shell.run_code("first.square()")
        with self.assertRaises(AttributeError):
            self.shell.run_code("first.cube()")
        with self.assertRaises(AttributeError):
            self.shell.run_code("first.power(5)")
        self.shell.run_code("first.b")
        with self.assertRaises(AttributeError):
            self.shell.run_code("first.toto")

        # remove square, add power

        self.write_file(
            mod_fn,
            textwrap.dedent(
                """
                            class MyClass:

                                def __init__(self, a=10):
                                    self.a = a
                                    self.b = 11

                                def power(self, p):
                                    print('compute power '+str(p))
                                    return self.a**p
                            """
            ),
        )

        self.shell.run_code("second = MyClass(5)")

        for object_name in {"first", "second"}:
            self.shell.run_code(f"{object_name}.power(5)")
            with self.assertRaises(AttributeError):
                self.shell.run_code(f"{object_name}.cube()")
            with self.assertRaises(AttributeError):
                self.shell.run_code(f"{object_name}.square()")
            self.shell.run_code(f"{object_name}.b")
            self.shell.run_code(f"{object_name}.a")
            with self.assertRaises(AttributeError):
                self.shell.run_code(f"{object_name}.toto")

    @skipif_not_numpy
    def test_comparing_numpy_structures(self):
        self.shell.magic_autoreload("2")
        self.shell.run_code("1+1")
        mod_name, mod_fn = self.new_module(
            textwrap.dedent(
                """
                                import numpy as np
                                class MyClass:
                                    a = (np.array((.1, .2)),
                                         np.array((.2, .3)))
                            """
            )
        )
        self.shell.run_code("from %s import MyClass" % mod_name)
        self.shell.run_code("first = MyClass()")

        # change property `a`
        self.write_file(
            mod_fn,
            textwrap.dedent(
                """
                                import numpy as np
                                class MyClass:
                                    a = (np.array((.3, .4)),
                                         np.array((.5, .6)))
                            """
            ),
        )

        with tt.AssertNotPrints(
            ("[autoreload of %s failed:" % mod_name), channel="stderr"
        ):
            self.shell.run_code("pass")  # trigger another reload

    def test_autoload_newly_added_objects(self):
        # All of these fail with %autoreload 2
        self.shell.magic_autoreload("3")
        mod_code = """
        def func1(): pass
        """
        mod_name, mod_fn = self.new_module(textwrap.dedent(mod_code))
        self.shell.run_code(f"from {mod_name} import *")
        self.shell.run_code("func1()")
        with self.assertRaises(NameError):
            self.shell.run_code("func2()")
        with self.assertRaises(NameError):
            self.shell.run_code("t = Test()")
        with self.assertRaises(NameError):
            self.shell.run_code("number")

        # ----------- TEST NEW OBJ LOADED --------------------------

        new_code = """
        def func1(): pass
        def func2(): pass
        class Test: pass
        number = 0
        from enum import Enum
        class TestEnum(Enum):
            A = 'a'
        """
        self.write_file(mod_fn, textwrap.dedent(new_code))

        # test function now exists in shell's namespace namespace
        self.shell.run_code("func2()")
        # test function now exists in module's dict
        self.shell.run_code(f"import sys; sys.modules['{mod_name}'].func2()")
        # test class now exists
        self.shell.run_code("t = Test()")
        # test global built-in var now exists
        self.shell.run_code("number")
        # test the enumerations gets loaded successfully
        self.shell.run_code("TestEnum.A")

        # ----------- TEST NEW OBJ CAN BE CHANGED --------------------

        new_code = """
        def func1(): return 'changed'
        def func2(): return 'changed'
        class Test:
            def new_func(self):
                return 'changed'
        number = 1
        from enum import Enum
        class TestEnum(Enum):
            A = 'a'
            B = 'added'
        """
        self.write_file(mod_fn, textwrap.dedent(new_code))
        self.shell.run_code("assert func1() == 'changed'")
        self.shell.run_code("assert func2() == 'changed'")
        self.shell.run_code("t = Test(); assert t.new_func() == 'changed'")
        self.shell.run_code("assert number == 1")
        if sys.version_info < (3, 12):
            self.shell.run_code("assert TestEnum.B.value == 'added'")

        # ----------- TEST IMPORT FROM MODULE --------------------------

        new_mod_code = """
        from enum import Enum
        class Ext(Enum):
            A = 'ext'
        def ext_func():
            return 'ext'
        class ExtTest:
            def meth(self):
                return 'ext'
        ext_int = 2
        """
        new_mod_name, new_mod_fn = self.new_module(textwrap.dedent(new_mod_code))
        current_mod_code = f"""
        from {new_mod_name} import *
        """
        self.write_file(mod_fn, textwrap.dedent(current_mod_code))
        self.shell.run_code("assert Ext.A.value == 'ext'")
        self.shell.run_code("assert ext_func() == 'ext'")
        self.shell.run_code("t = ExtTest(); assert t.meth() == 'ext'")
        self.shell.run_code("assert ext_int == 2")

    def test_verbose_names(self):
        # Asserts correspondense between original mode names and their verbose equivalents.
        @dataclass
        class AutoreloadSettings:
            check_all: bool
            enabled: bool
            autoload_obj: bool

        def gather_settings(mode):
            self.shell.magic_autoreload(mode)
            module_reloader = self.shell.auto_magics._reloader
            return AutoreloadSettings(
                module_reloader.check_all,
                module_reloader.enabled,
                module_reloader.autoload_obj,
            )

        assert gather_settings("0") == gather_settings("off")
        assert gather_settings("0") == gather_settings("OFF")  # Case insensitive
        assert gather_settings("1") == gather_settings("explicit")
        assert gather_settings("2") == gather_settings("all")
        assert gather_settings("3") == gather_settings("complete")

        # And an invalid mode name raises an exception.
        with self.assertRaises(ValueError):
            self.shell.magic_autoreload("4")

    def test_aimport_parsing(self):
        # Modules can be included or excluded all in one line.
        module_reloader = self.shell.auto_magics._reloader
        self.shell.magic_aimport("os")  # import and mark `os` for auto-reload.
        assert module_reloader.modules["os"] is True
        assert "os" not in module_reloader.skip_modules.keys()

        self.shell.magic_aimport("-math")  # forbid autoreloading of `math`
        assert module_reloader.skip_modules["math"] is True
        assert "math" not in module_reloader.modules.keys()

        self.shell.magic_aimport(
            "-os, math"
        )  # Can do this all in one line; wasn't possible before.
        assert module_reloader.modules["math"] is True
        assert "math" not in module_reloader.skip_modules.keys()
        assert module_reloader.skip_modules["os"] is True
        assert "os" not in module_reloader.modules.keys()

    def test_autoreload_output(self):
        self.shell.magic_autoreload("complete")
        mod_code = """
        def func1(): pass
        """
        mod_name, mod_fn = self.new_module(mod_code)
        self.shell.run_code(f"import {mod_name}")
        with tt.AssertPrints("", channel="stdout"):  # no output; this is default
            self.shell.run_code("pass")

        self.shell.magic_autoreload("complete --print")
        self.write_file(mod_fn, mod_code)  # "modify" the module
        with tt.AssertPrints(
            f"Reloading '{mod_name}'.", channel="stdout"
        ):  # see something printed out
            self.shell.run_code("pass")

        self.shell.magic_autoreload("complete -p")
        self.write_file(mod_fn, mod_code)  # "modify" the module
        with tt.AssertPrints(
            f"Reloading '{mod_name}'.", channel="stdout"
        ):  # see something printed out
            self.shell.run_code("pass")

        self.shell.magic_autoreload("complete --print --log")
        self.write_file(mod_fn, mod_code)  # "modify" the module
        with tt.AssertPrints(
            f"Reloading '{mod_name}'.", channel="stdout"
        ):  # see something printed out
            self.shell.run_code("pass")

        self.shell.magic_autoreload("complete --print --log")
        self.write_file(mod_fn, mod_code)  # "modify" the module
        with self.assertLogs(logger="autoreload") as lo:  # see something printed out
            self.shell.run_code("pass")
        assert lo.output == [f"INFO:autoreload:Reloading '{mod_name}'."]

        self.shell.magic_autoreload("complete -l")
        self.write_file(mod_fn, mod_code)  # "modify" the module
        with self.assertLogs(logger="autoreload") as lo:  # see something printed out
            self.shell.run_code("pass")
        assert lo.output == [f"INFO:autoreload:Reloading '{mod_name}'."]

    def _check_smoketest(self, use_aimport=True):
        """
        Functional test for the automatic reloader using either
        '%autoreload 1' or '%autoreload 2'
        """

        mod_name, mod_fn = self.new_module(
            """
x = 9

z = 123  # this item will be deleted

def foo(y):
    return y + 3

class Baz(object):
    def __init__(self, x):
        self.x = x
    def bar(self, y):
        return self.x + y
    @property
    def quux(self):
        return 42
    def zzz(self):
        '''This method will be deleted below'''
        return 99

class Bar:    # old-style class: weakref doesn't work for it on Python < 2.7
    def foo(self):
        return 1
"""
        )

        #
        # Import module, and mark for reloading
        #
        if use_aimport:
            self.shell.magic_autoreload("1")
            self.shell.magic_aimport(mod_name)
            stream = StringIO()
            self.shell.magic_aimport("", stream=stream)
            self.assertIn(("Modules to reload:\n%s" % mod_name), stream.getvalue())

            with self.assertRaises(ImportError):
                self.shell.magic_aimport("tmpmod_as318989e89ds")
        else:
            self.shell.magic_autoreload("2")
            self.shell.run_code("import %s" % mod_name)
            stream = StringIO()
            self.shell.magic_aimport("", stream=stream)
            self.assertTrue(
                "Modules to reload:\nall-except-skipped" in stream.getvalue()
            )
        self.assertIn(mod_name, self.shell.ns)

        mod = sys.modules[mod_name]

        #
        # Test module contents
        #
        old_foo = mod.foo
        old_obj = mod.Baz(9)
        old_obj2 = mod.Bar()

        def check_module_contents():
            self.assertEqual(mod.x, 9)
            self.assertEqual(mod.z, 123)

            self.assertEqual(old_foo(0), 3)
            self.assertEqual(mod.foo(0), 3)

            obj = mod.Baz(9)
            self.assertEqual(old_obj.bar(1), 10)
            self.assertEqual(obj.bar(1), 10)
            self.assertEqual(obj.quux, 42)
            self.assertEqual(obj.zzz(), 99)

            obj2 = mod.Bar()
            self.assertEqual(old_obj2.foo(), 1)
            self.assertEqual(obj2.foo(), 1)

        check_module_contents()

        #
        # Simulate a failed reload: no reload should occur and exactly
        # one error message should be printed
        #
        self.write_file(
            mod_fn,
            """
a syntax error
""",
        )

        with tt.AssertPrints(
            ("[autoreload of %s failed:" % mod_name), channel="stderr"
        ):
            self.shell.run_code("pass")  # trigger reload
        with tt.AssertNotPrints(
            ("[autoreload of %s failed:" % mod_name), channel="stderr"
        ):
            self.shell.run_code("pass")  # trigger another reload
        check_module_contents()

        #
        # Rewrite module (this time reload should succeed)
        #
        self.write_file(
            mod_fn,
            """
x = 10

def foo(y):
    return y + 4

class Baz(object):
    def __init__(self, x):
        self.x = x
    def bar(self, y):
        return self.x + y + 1
    @property
    def quux(self):
        return 43

class Bar:    # old-style class
    def foo(self):
        return 2
""",
        )

        def check_module_contents():
            self.assertEqual(mod.x, 10)
            self.assertFalse(hasattr(mod, "z"))

            self.assertEqual(old_foo(0), 4)  # superreload magic!
            self.assertEqual(mod.foo(0), 4)

            obj = mod.Baz(9)
            self.assertEqual(old_obj.bar(1), 11)  # superreload magic!
            self.assertEqual(obj.bar(1), 11)

            self.assertEqual(old_obj.quux, 43)
            self.assertEqual(obj.quux, 43)

            self.assertFalse(hasattr(old_obj, "zzz"))
            self.assertFalse(hasattr(obj, "zzz"))

            obj2 = mod.Bar()
            self.assertEqual(old_obj2.foo(), 2)
            self.assertEqual(obj2.foo(), 2)

        self.shell.run_code("pass")  # trigger reload
        check_module_contents()

        #
        # Another failure case: deleted file (shouldn't reload)
        #
        os.unlink(mod_fn)

        self.shell.run_code("pass")  # trigger reload
        check_module_contents()

        #
        # Disable autoreload and rewrite module: no reload should occur
        #
        if use_aimport:
            self.shell.magic_aimport("-" + mod_name)
            stream = StringIO()
            self.shell.magic_aimport("", stream=stream)
            self.assertTrue(("Modules to skip:\n%s" % mod_name) in stream.getvalue())

            # This should succeed, although no such module exists
            self.shell.magic_aimport("-tmpmod_as318989e89ds")
        else:
            self.shell.magic_autoreload("0")

        self.write_file(
            mod_fn,
            """
x = -99
""",
        )

        self.shell.run_code("pass")  # trigger reload
        self.shell.run_code("pass")
        check_module_contents()

        #
        # Re-enable autoreload: reload should now occur
        #
        if use_aimport:
            self.shell.magic_aimport(mod_name)
        else:
            self.shell.magic_autoreload("")

        self.shell.run_code("pass")  # trigger reload
        self.assertEqual(mod.x, -99)

    def test_smoketest_aimport(self):
        self._check_smoketest(use_aimport=True)

    def test_smoketest_autoreload(self):
        self._check_smoketest(use_aimport=False)
```
