## Supplementary 2 for "A Universal RT-qPCR Method for DNA Aptamer Quantification": test_storemagic.py

```
import tempfile, os
from pathlib import Path

from traitlets.config.loader import Config


def setup_module():
    ip.magic('load_ext storemagic')

def test_store_restore():
    assert 'bar' not in ip.user_ns, "Error: some other test leaked `bar` in user_ns"
    assert 'foo' not in ip.user_ns, "Error: some other test leaked `foo` in user_ns"
    assert 'foobar' not in ip.user_ns, "Error: some other test leaked `foobar` in user_ns"
    assert 'foobaz' not in ip.user_ns, "Error: some other test leaked `foobaz` in user_ns"
    ip.user_ns['foo'] = 78
    ip.magic('alias bar echo "hello"')
    ip.user_ns['foobar'] = 79
    ip.user_ns['foobaz'] = '80'
    tmpd = tempfile.mkdtemp()
    ip.magic('cd ' + tmpd)
    ip.magic('store foo')
    ip.magic('store bar')
    ip.magic('store foobar foobaz')

    # Check storing
    assert ip.db["autorestore/foo"] == 78
    assert "bar" in ip.db["stored_aliases"]
    assert ip.db["autorestore/foobar"] == 79
    assert ip.db["autorestore/foobaz"] == "80"

    # Remove those items
    ip.user_ns.pop('foo', None)
    ip.user_ns.pop('foobar', None)
    ip.user_ns.pop('foobaz', None)
    ip.alias_manager.undefine_alias('bar')
    ip.magic('cd -')
    ip.user_ns['_dh'][:] = []

    # Check restoring
    ip.magic("store -r foo bar foobar foobaz")
    assert ip.user_ns["foo"] == 78
    assert ip.alias_manager.is_alias("bar")
    assert ip.user_ns["foobar"] == 79
    assert ip.user_ns["foobaz"] == "80"

    ip.magic("store -r")  # restores _dh too
    assert any(Path(tmpd).samefile(p) for p in ip.user_ns["_dh"])

    os.rmdir(tmpd)

def test_autorestore():
    ip.user_ns['foo'] = 95
    ip.magic('store foo')
    del ip.user_ns['foo']
    c = Config()
    c.StoreMagics.autorestore = False
    orig_config = ip.config
    try:
        ip.config = c
        ip.extension_manager.reload_extension("storemagic")
        assert "foo" not in ip.user_ns
        c.StoreMagics.autorestore = True
        ip.extension_manager.reload_extension("storemagic")
        assert ip.user_ns["foo"] == 95
    finally:
        ip.config = orig_config
```
