## Supplementary 2 for "A Universal RT-qPCR Method for DNA Aptamer Quantification": README.txt

=======================================================
 Nose plugin with IPython and extension module support
=======================================================

This directory provides the key functionality for test support that IPython
needs as a nose plugin, which can be installed for use in projects other than
IPython.

The presence of a Makefile here is mostly for development and debugging
purposes as it only provides a few shorthand commands.  You can manually
install the plugin by using standard Python procedures (``setup.py install``
with appropriate arguments).

To install the plugin using the Makefile, edit its first line to reflect where
you'd like the installation.

Once you've set the prefix, simply build/install the plugin with::

  make

and run the tests with::

  make test

You should see output similar to::

  maqroll[plugin]> make test
  nosetests -s --with-ipdoctest --doctest-tests dtexample.py
  ..
  ----------------------------------------------------------------------
  Ran 2 tests in 0.016s

  OK
