## Supplementary 2 for "A Universal RT-qPCR Method for DNA Aptamer Quantification": test_combo.txt

=======================
 Combo testing example
=======================

This is a simple example that mixes ipython doctests::

    In [1]: import code

    In [2]: 2**12
    Out[2]: 4096

with command-line example information that does *not* get executed::

    $ mpirun -n 4 ipengine --controller-port=10000 --controller-ip=host0

and with literal examples of Python source code::

    controller = dict(host='myhost',
		      engine_port=None, # default is 10105
		      control_port=None,
		      )

    # keys are hostnames, values are the number of engine on that host
    engines = dict(node1=2,
		   node2=2,
		   node3=2,
		   node3=2,
		   )

    # Force failure to detect that this test is being run.
    1/0

These source code examples are executed but no output is compared at all.  An
error or failure is reported only if an exception is raised.

NOTE: the execution of pure python blocks is not yet working!
