## Supplementary 2 for "A Universal RT-qPCR Method for DNA Aptamer Quantification": test_example.txt

=====================================
 Tests in example form - pure python
=====================================

This file contains doctest examples embedded as code blocks, using normal
Python prompts.  See the accompanying file for similar examples using IPython
prompts (you can't mix both types within one file). The following will be run
as a test::

    >>> 1+1
    2
    >>> print ("hello")
    hello

More than one example works::

    >>> s="Hello World"

    >>> s.upper()
    'HELLO WORLD'

but you should note that the *entire* test file is considered to be a single
test.  Individual code blocks that fail are printed separately as ``example
failures``, but the whole file is still counted and reported as one test.
