## Supplementary 2 for "A Universal RT-qPCR Method for DNA Aptamer Quantification": test_exampleip.txt

=================================
 Tests in example form - IPython
=================================

You can write text files with examples that use IPython prompts (as long as you
use the nose ipython doctest plugin), but you can not mix and match prompt
styles in a single file.  That is, you either use all ``>>>`` prompts or all
IPython-style prompts.  Your test suite *can* have both types, you just need to
put each type of example in a separate. Using IPython prompts, you can paste
directly from your session::

    In [5]: s="Hello World"

    In [6]: s.upper()
    Out[6]: 'HELLO WORLD'

Another example::

    In [8]: 1+3
    Out[8]: 4

Just like in IPython docstrings, you can use all IPython syntax and features::

    In [9]: !echo hello
    hello

    In [10]: a='hi'

    In [11]: !echo $a
    hi
