## Supplementary 2 for "A Universal RT-qPCR Method for DNA Aptamer Quantification": Aptamer Primer Designer Manual.docx

### 1. Introduction

Aptamers are generally too short to allow direct identification of suitable primer pairs. Therefore, primers need to be generated de novo using the software “Aptamer Primer Designer” for aptamer RT-qPCR. Our approach involves generating a large number of random primers, which are then filtered to identify those that match the aptamer sequence.

Briefly, random primer pools are generated and then filtered to match the aptamer sequence. The design process differs slightly depending on the aptamer type—whether it contains one primer site or none. For aptamers with one primer site, the software identifies the existing site, and a second primer is selected from the primer pool. For aptamers without a primer site, primer pairs are first generated from the pool and then screened against the aptamer sequence to identify suitable combinations. Probe design follows a similar strategy.

The overall workflow is illustrated in Figure 1. You should design primers sequentially using tools 2 through 5, followed by designing probes using tools 6 through 8.

Note: Most parameters can be adjusted according to your experimental environment or specific needs. The default parameter set provided has been validated and shown to perform effectively in the current study.

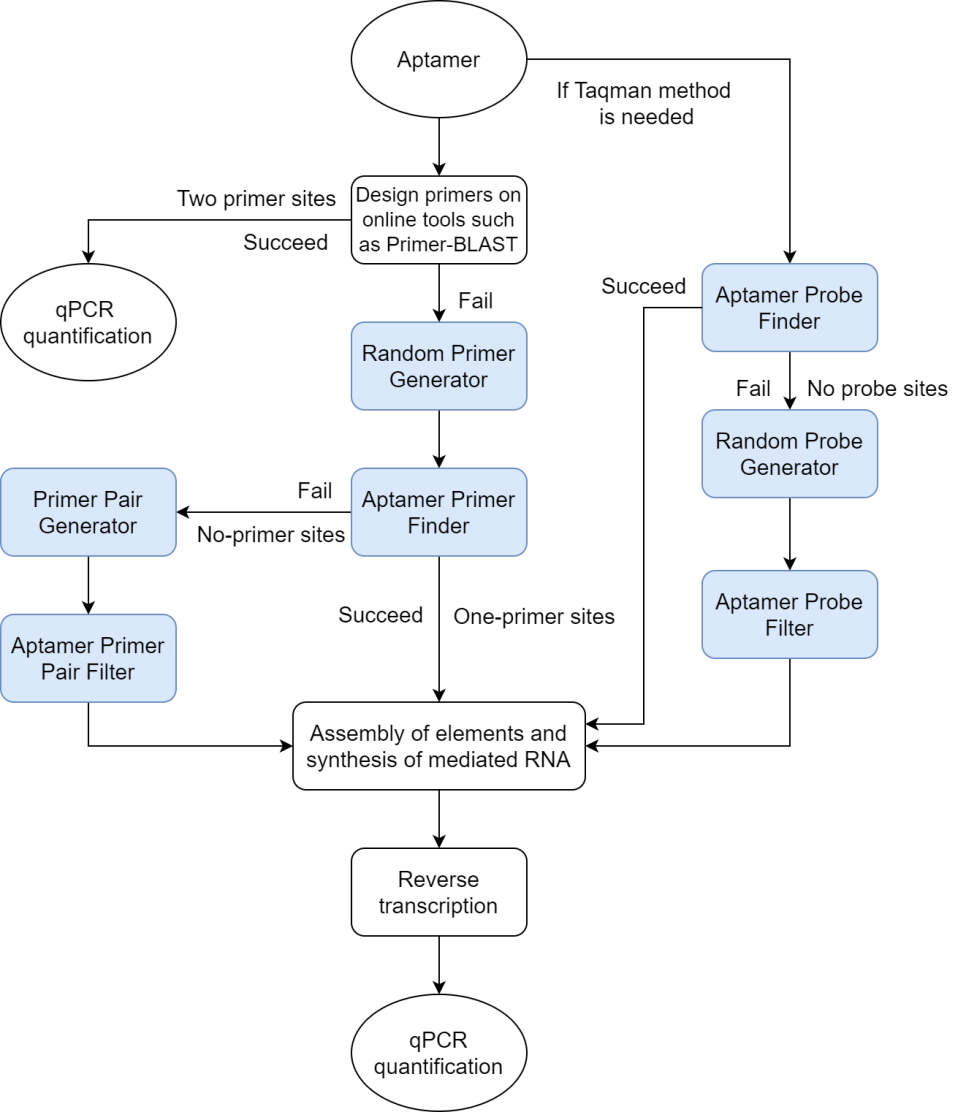

Figure 1 Flowchart for using “Aptamer Primer Designer” software.

### 2. Random Primer Generator

Random primers with varying melting temperatures (Tm) are generated through a random mutation algorithm, adhering to typical primer design guidelines.

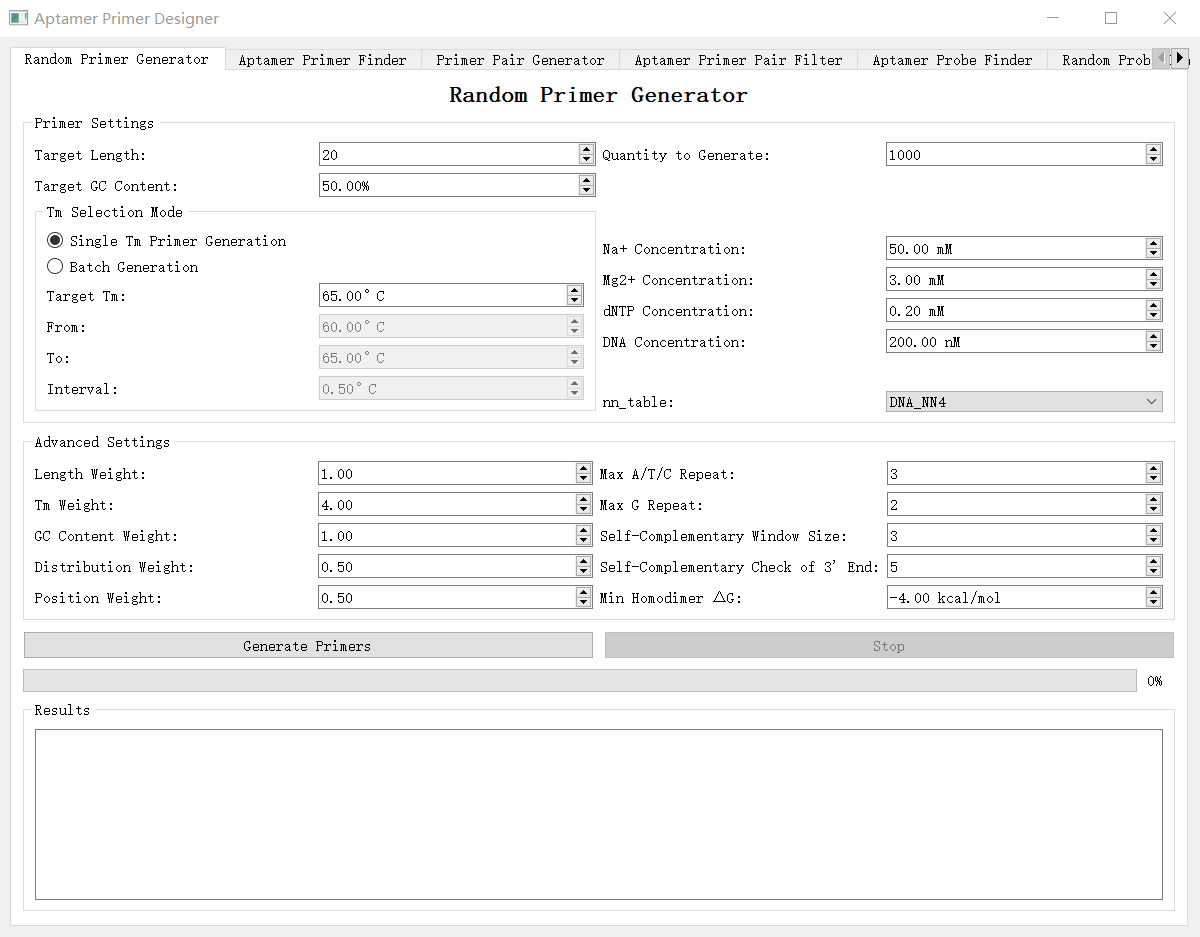

Figure 2.1 User interface (UI) of the "Random Primer Generator" module.

A. Generation Mode

**Single Tm Primer Generation:** Generates a single Tm-specific primer pool.

**Batch Generation:** Generates multiple primer pools within a specified Tm range. For example, setting the range "60 to 65, interval 0.5" will generate primer pools with Tm values of 60, 60.5, 61, 61.5, ..., 65℃. All files will be generated simultaneously.

B. Basic parameters

**Tm (℃)**: Generally within the range of 60-65°C.

**Primer Length**: Typically between 18-25 nucleotides.

**GC Content (%)**: Aiming for approximately 50%.

**Quantity**: Typically, a pool of 1000 primers is sufficient for screening purposes.

**Weight Setting**: The algorithm generates primer sequences based on five parameters: **Length, Tm, GC content, base distribution,** and **base position**. As the weight of a parameter increases, the algorithm gives it more consideration during primer generation. For example, increasing the weight of base distribution balances the proportions of A, T, C, and G, while increasing the weight of base position results in a more uniform distribution of these bases across the primer.

C. qPCR environmental variables

Set the following parameters based on your specific qPCR conditions:

**Monovalent Cation Concentration (mM):** The concentration of monovalent cations, such as Na^+^.

**Divalent Cation Concentration (mM):** The concentration of divalent cations, such as Mg^2+^.

**dNTP Concentration (mM):** The concentration of dNTP.

**DNA Concentration (nM):** The concentration of one of primers in qPCR.

D. Algorithm

**nn_table**: The algorithm utilizes four methods for calculating DNA/DNA Tm, as implemented in the Python package “Bio.SeqUtils” ([https://biopython.org/docs/1.75/api/Bio.SeqUtils.MeltingTemp.html](https://biopython.org/docs/1.75/api/Bio.SeqUtils.MeltingTemp.html" \t "_new), this resource offers a detailed explanation of the methods). Among them, **mt.DNA_NN4** is the newest approach and has been evaluated in the present study. Different primer design tools may employ varying algorithms; thus, you can select an alternative method if compatibility with other tools is needed.

E. Primer check parameters

**Max A/T/C Repeat**: Ideally, fewer than 4 consecutive occurrences are preferred; for example, “AAAA” is avoided.

**Max G Repeat**: Ideally, fewer than 3 consecutive “G” are preferred.

**Self-Complementary Window Size**: The number of bases in a primer that complement to each other (Figure 2.2). A higher count increases the risk of primer dimer formation, but overly strict constraints can significantly extend the program's runtime.

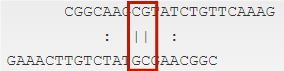

Figure 2.2 An example of “window” (red, window size 2).

**Self-Complementary Check of 3’ End**: The primer’s 3’ end should theoretically avoid self-complementarity. In this study, we define this rule as "the last X bases should not contain a complementary window of 2 bases." Increasing X reduces the likelihood of dimer formation. Our tests indicate that a value of 5 is effective, though other values remain untested.

**Min Homodimer ΔG (kcal/mol)**: **Δ**G represents the Gibbs free energy change during homodimer formation, indicating the stability of the interaction. A lower **Δ**G value suggests a higher likelihood of dimer formation. Generally, **Δ**G should be greater than -6 to minimize dimerization.

These parameters are essential for generating high-quality primers. However, more stringent parameter settings may reduce the likelihood of generating viable primers and can significantly increase program runtime. The default settings are balanced to optimize both time efficiency and primer quality based on our test conditions.

F. Other settings

**Generate Primers/Stop**: To start or stop the program. Stopping the program does not delete existing data. If you restart, newly generated primers will be appended to the existing data unless you have manually removed the corresponding files.

Generated primer pools are automatically saved in the program folder with the filename format “primers+Tm”.

### 3. Aptamer Primer Finder

Detect if the aptamer has single available primer sites.

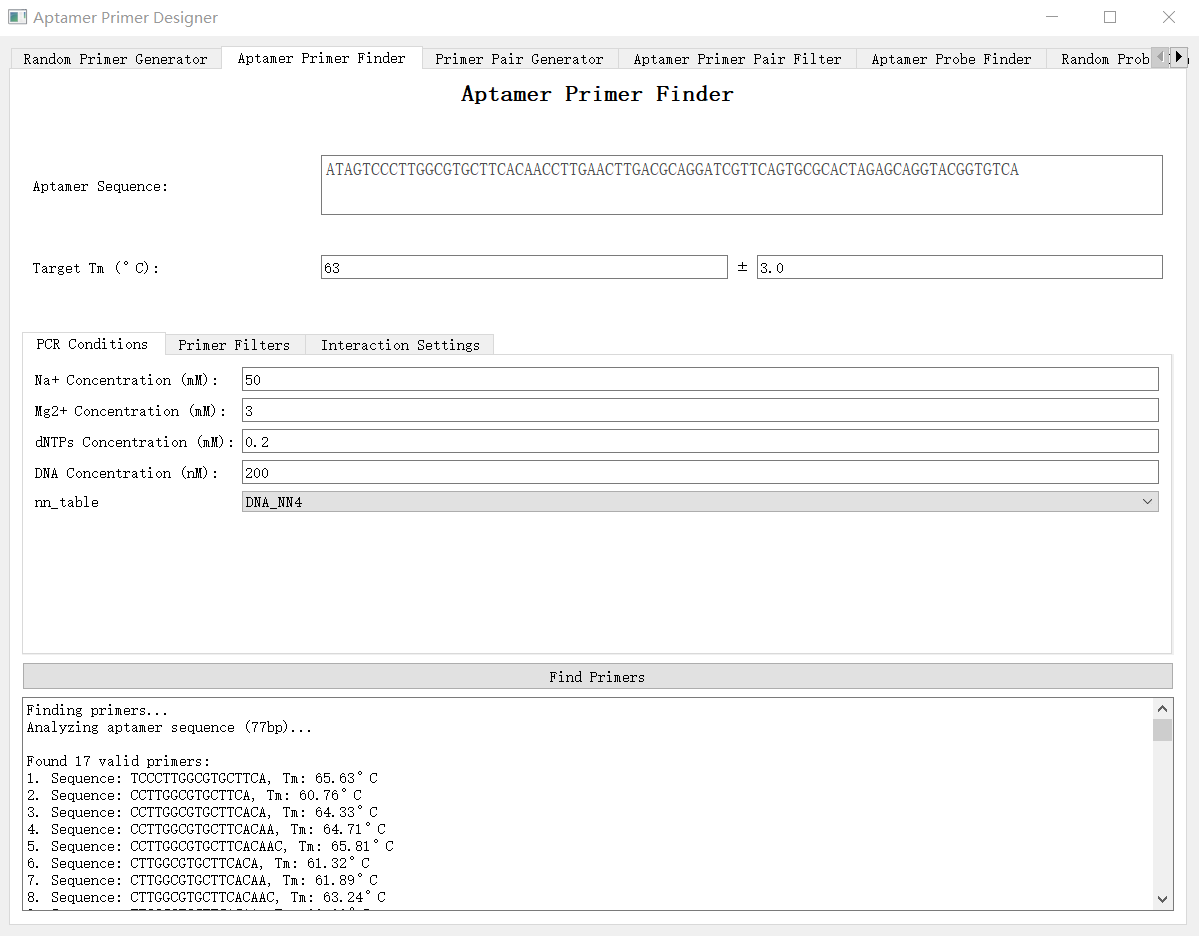

Figure 3 UI of the "Aptamer Primer Finder" module.

A. Basic Parameters

**Apt Sequence:** Input the aptamer sequence.

**Tm Range (℃):** Set the primer melting temperature (Tm) range.

B. qPCR environmental variables and Algorithm

Same as in the "Random Primer Generator".

C. Primer check parameters

**Min/Max GC Content (%):** Sets the GC content range, e.g., 40-60%.

**Min/Max C/G at Last 5 Bases:** Defines the allowable GC count within the last 5 bases, typically 1 or 2.

Other parameters are same as in the "Random Primer Generator".

D. Primer pairs check parameters

Primers from random primer pools are filtered based on the following rules:

**Aptamer Interaction Window Size:** Window size of complement between aptamer and primer from pool.

**Primer Pair Complementary Window Size:** Number of complementary bases between primer pairs.

**Primer Pair Complementary Check of 3' End :** The last X bases of the primer pair must not contain a complementary window of length Y (**3’ End Complementary Check Window Size**).

Once you click "Find Primers," the module will search for suitable primer sites on the aptamer (Figure 3.1) and automatically screen the primer pools to generate optimal primer pairs. Results will be displayed on the panel, and you can adjust parameters based on this feedback.

Total primers passing Tm filter: The total number of primers identified within the specified Tm range.

CG end exclusions: Primers excluded due to **Min/Max C/G at Last 5 Bases**.

Single base repeat exclusions: Primers excluded due to **Max A/T/C Repeat.**

Consecutive G exclusions: Primers excluded due to **Max G Repeat.**

Reverse complement exclusions: Primers excluded due to **Self-Complementary Window Size.**

3' end exclusions: Primers excluded due to **Self-Complementary Check of 3’ End.**

Homodimer ΔG exclusions: Primers excluded due to **Min Homodimer ΔG (kcal/mol)**.

GC content exclusions: Primers excluded due to **Min/Max GC Content (%)**.

- Excluded X primer2 candidates due to aptamer interaction: Primers excluded due to **Aptamer Interaction Window Size**.

- Excluded X primer2 candidates due to primer-primer interaction: Primers excluded due to **Primer Pair Complementary Window Size** and **Primer Pair Complementary Check of 3' End**.

The generated primer pairs results will be auto-saved to 'Valid_Primer_Pairs.csv'.

### 4. Primer Pair Generator

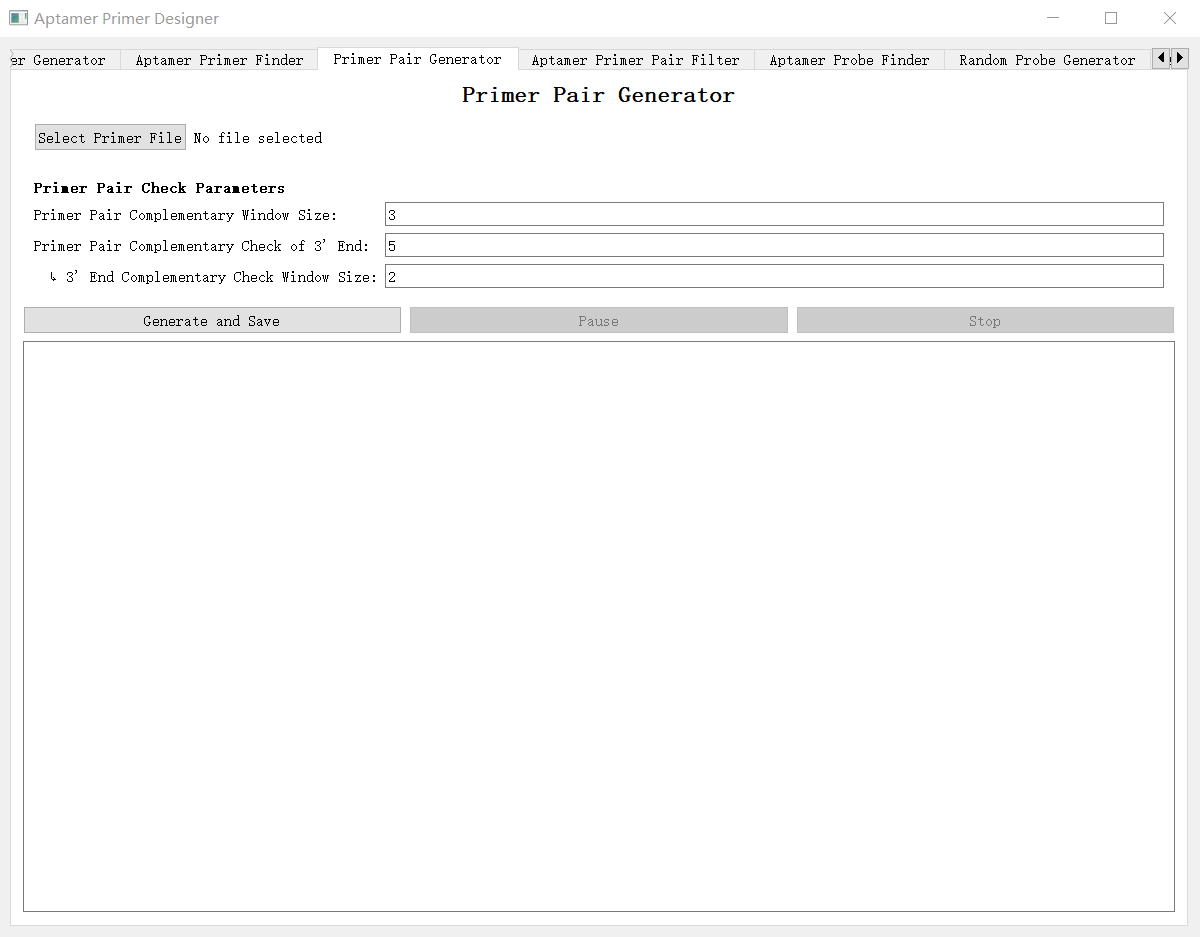

Figure 4 UI of the "Primer Pair Generator" module.

This module generates primer pairs from selected primer pools. Begin by selecting a primer pool file. Primer pairs will be screened based on their interactions, using the same constraint parameters described previously. The generated primer pairs pool will be auto-saved to 'primer_pairs_of_primers + Tm.csv'.

### 5. Aptamer Primer Pair Filter

Filter primer pairs from the primer pair pool to ensure compatibility with the aptamer sequence.

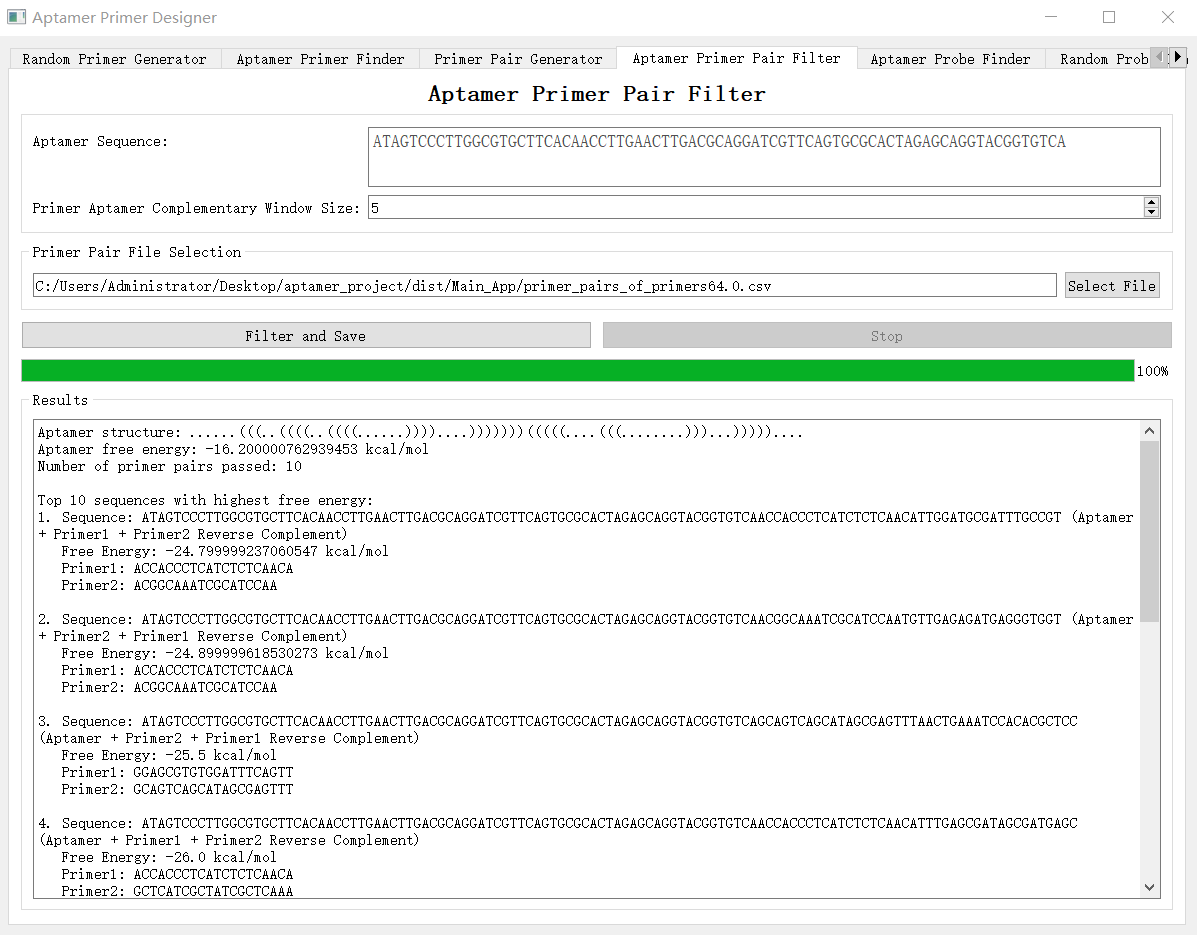

Figure 5 UI of the "Aptamer Primer Pair Filter" module.

**Apt Sequence:** Input the aptamer sequence.

**Primer Aptamer Complementary Window Size:** The complementary window between each primer and aptamer. Both primers should pass this check.
**Primer Pair File Selection:** Select a primer pair pool to filter and identify available primer pairs.

**Filter and Save:** Initiate the filtering process and save the results to a document. Filtered primer pairs will be ranked according to the free energy of the combined [Aptamer + Primers] sequence, with higher values prioritized to minimize potential secondary structures.

### 6. Apt Probe Finder

Identify available probes within the aptamer sequence.

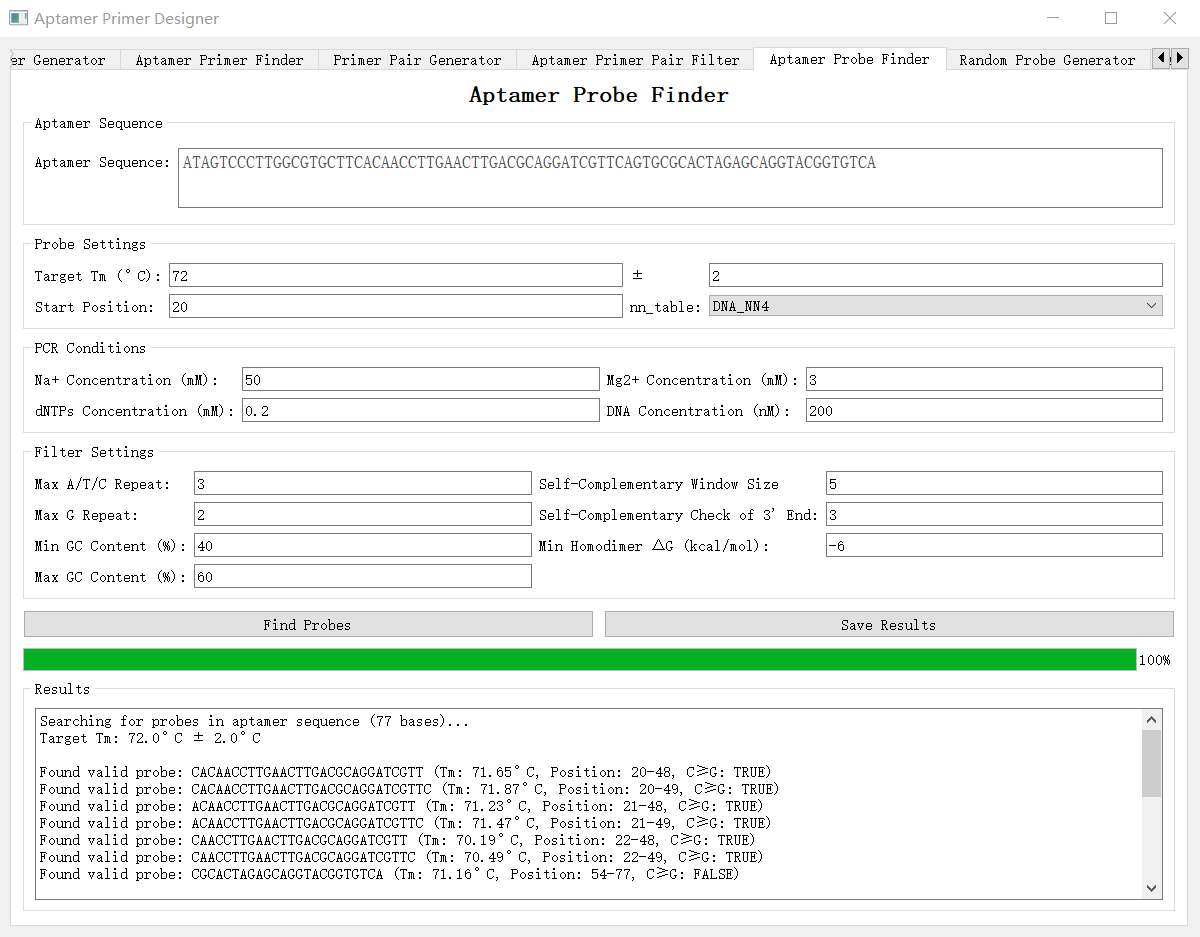

Figure 6 UI of the "Apt Probe Finder" module.

**Probe Length:** Set the target length of the generated probes.

**Start Position:** Define the starting position for probe selection. For example, for the sequence AATCGGA, if you want to start probe filtering from C (assuming AAT is a primer site), the start position should be 4.

**Note:** If C < G in the probe sequence, synthesize the reverse complementary sequence as the probe.

### 7. Random Probe Generator

Generates a random probe pool with a specified Tm.

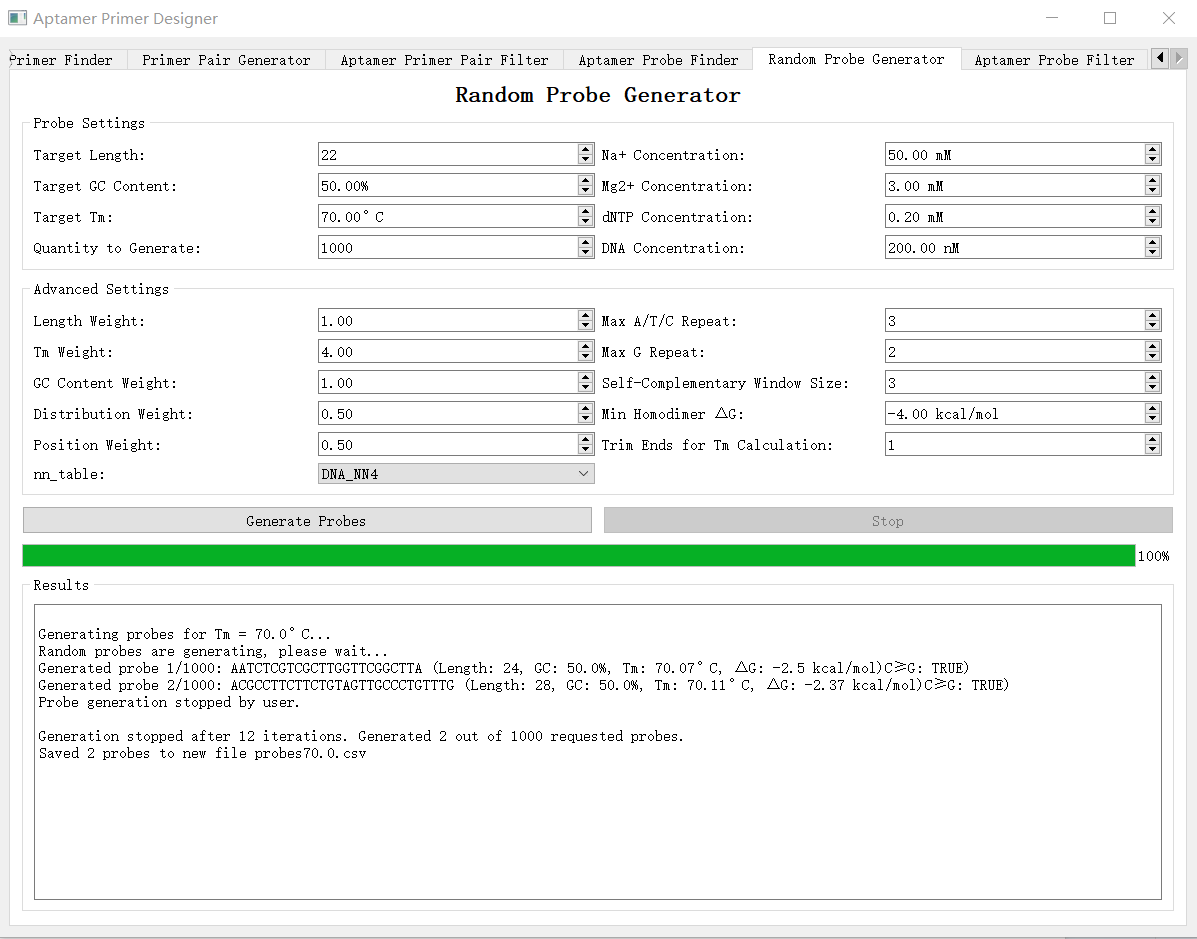

Figure 7 UI of the "Random Probe Generator" module.

**Trim Ends for Tm Calculation:** In certain cases, an interval between primer and probe sites is needed. You can set this interval size to specify the number of nucleotides to exclude from Tm calculation.

Example:
Consider the sequence CTCTAGAG, where AGAG is the primer site, and you intend to insert a probe immediately upstream, leaving a 2-base interval between the probe and primer. If you set the interval size to 2, the tool will generate a probe like XXAGCGTXX, but it will calculate the Tm only using the internal portion (AGCGT). Thus, your final assembled sequence will be CTCTXXAGCGTXXAGAG and probe is AGCGT.

### 8. Apt Probe Filter

Once a primer pair has been selected, if no suitable probe-binding site is available within the aptamer sequence, this module will filter available probes according to compatibility criteria.

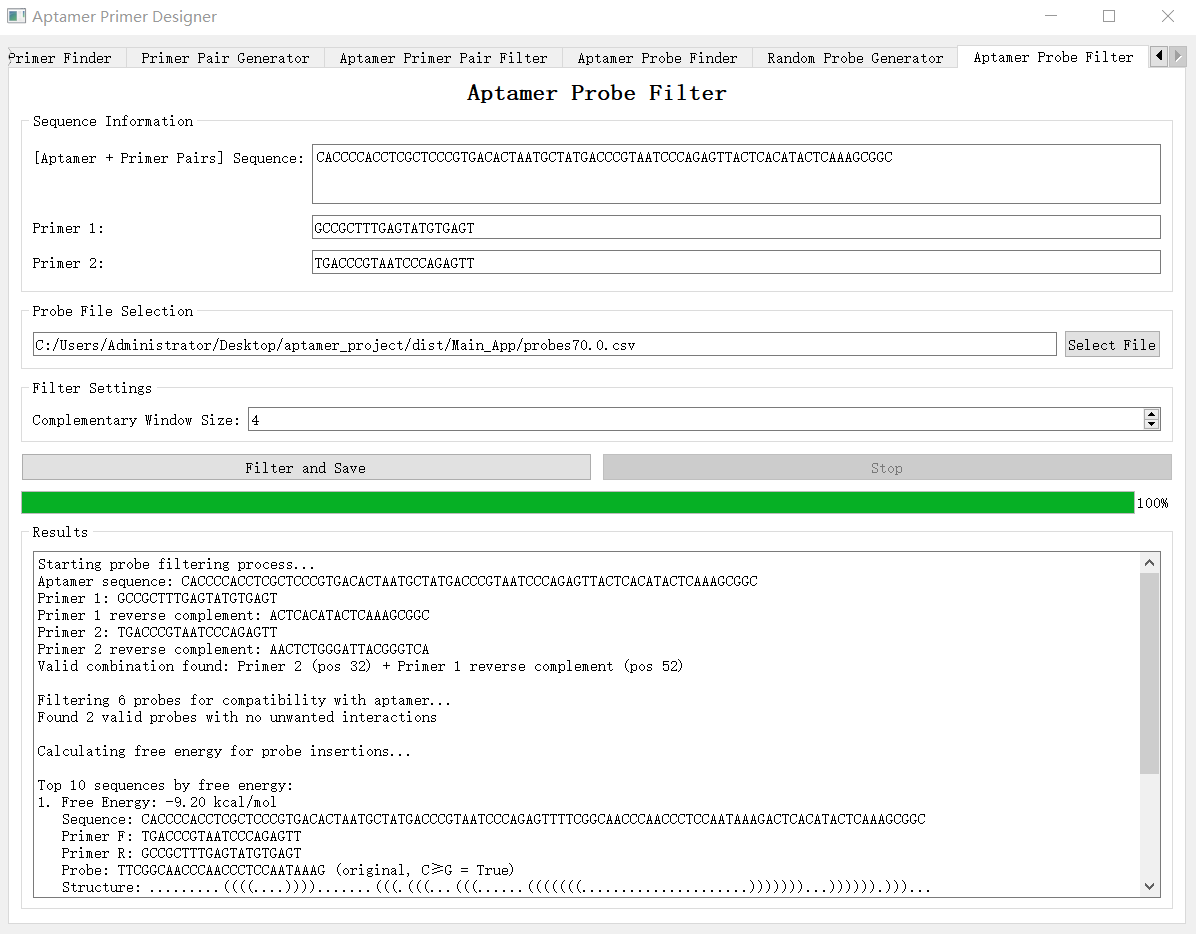

**[Aptamer + Primer Pairs] Sequence:** Paste the complete [Aptamer + Primer Pairs] sequence obtained from the "**Aptamer Primer Finder**" or "**Aptamer Primer Pair Filter**," consisting of Aptamer + Primer F + reverse complementary Primer R.

**Primer1:** Filtered or generated primer sequence.

**Primer2:** Generated primer sequence.

**Probe pool:** Set a probe pool to filter.

**Complementary Window Size:** Window size for checking complementarity within the [Aptamer + Primer Pairs] sequence.
